# ACME: an Affinity-based Cas9 Mediated Enrichment method for targeted nanopore sequencing

**DOI:** 10.1101/2022.02.03.478550

**Authors:** Shruti V Iyer, Melissa Kramer, Sara Goodwin, W. Richard McCombie

## Abstract

Targeted sequencing significantly improves accuracy and coverage and aids in providing the depth necessary to detect rare alleles in a heterogenous population of cells. Until the introduction of nanopore Cas9 Targeted-Sequencing (nCATS), a lack of efficient long-read compatible targeting techniques made it difficult to study specific regions of interest on long-read platforms. Existing nCATS-based strategies are currently limited by the per molecule target lengths capturable (<30kb), requiring several Cas9 guides to tile across larger regions of interest, ultimately reducing the number of targets that can be surveyed per reaction. Also, longer read lengths help reduce mapping errors, making it more likely that complex structural rearrangements can be resolved. Absence of a background reduction step in nCATS also increases the competition between non-target and target fragments in the sequencing pool for pore occupancy, decreasing the overall percentage of on-target reads. To address this, we introduce ACME - an <u>A</u>ffinity-based <u>C</u>as9-<u>M</u>ediated <u>E</u>nrichment method - that helps reduce background reads, increasing on-target coverage and size of target regions that can be spanned with single reads to 100kb.

ACME uses a HisTag-based isolation and pulldown of Cas-9 bound non-target reads, reducing the background noise in sequencing. We designed a panel of guide RNAs targeting 10 genes to enrich for specific regions of the cancer genome and tested them in two breast cell lines – MCF 10A and SK-BR-3. These gene targets spanned different size ranges (10kb to 150kb) allowing us to identify the largest target sizes that could be optimally captured by single molecules spanning the entire region. When compared with using just nCATS, the ACME method for background reduction increased the overall coverage across the entire length of all targets by 2-fold to 25-fold. By using ACME to eliminate smaller competing non-targets from the sequencing library, we saw a 3- to 7-fold increase in the number of reads spanning 100% of the gene targets when compared to nCATS. For one of our larger targets, *BRCA2*, we observed >60-fold target enrichment, close to 70x coverage, and 3-20 reads spanning the entire 95kb target. We observed an increase in enrichment, depth, and number of whole gene spanning reads for other genes on the panel as well across both cell lines, with enrichment as high as 4000-fold for some genes. Furthermore, ACME identified all SVs previously called within our targets by ONT and PacBio whole genome sequencing and performed on par with these platforms for SNP detection when compared with Illumina short-read whole genome sequencing.

## Introduction

Cancer genomes have largely been evaluated using short-read sequencing technologies, which lack sensitivity and exhibit very high false positive rates (up to 89%) in SV detection ^1–4^. Long-read sequencing methods such as single-molecule real-time sequencing (SMRT) ^5^ and nanopore sequencing ^6^ have the ability to detect SVs with >=95% sensitivity and specificity ^7–9^. However, the difficulty in generating high enough sequence depth to detect rare alleles in a heterogeneous cell mix is even more difficult than with short reads because of the higher sequencing cost associated with long-read technologies. Targeted sequencing dramatically changes our ability to study the genome by facilitating higher sample throughput than whole genome sequencing and improves accuracy by increasing the read depth coverage ^10^. With targeted long-read sequencing, the goal is to be able to cover an entire target with several single contiguous reads spanning the whole region. Reads covering whole target regions minimize mapping errors due to SVs, aiding in their detection.

A Cas9-based enrichment approach called nanopore Cas9 Targeted-Sequencing (nCATS) was adapted for the Oxford Nanopore Technologies (ONT) platform by Gilpatrick *et al*., with singleended cleavage variations developed by Stangl *et al*. ^11^ to detect fusion events and Watson *et al*. ^12^ to detect genomic duplications. While nCATS is a fast and effective long-read targeting approach, its two major limitations are: 1) Relatively shorter read lengths (<30kb) leading to few or no reads that span larger targets from end to end. This would require several crRNA guides to tile through larger target regions, thereby limiting the number of targets that can be investigated in a single run. 2) Lack of an efficient background reduction step. The nCATS sequencing pool consists of target DNA fragments along with Cas9-bound and non-Cas9 bound dephosphorylated fragments. Higher ratio of non-target to target DNA in the prepared library often results in shorter non-target fragments competing with long targets for pore occupancy, effectively reducing total target depth.

Two groups have adapted the principle behind nCATS to perform exonuclease digestion of non-target DNA. Cas9-based Background Elimination (CaBagE) ^13^ and Negative Enrichment ^14^ focus on protecting target DNA ends with Cas9, followed by exonuclease digestion of non-target reads. However, both methods have only achieved target read lengths of <35kb. It is also important to note that this approach poses an additional risk of digesting target fragments that may have breaks between the Cas9 binding sites and are not protected by Cas9. Despite its potential, this approach failed to outperform nCATS in its yield and target coverage further highlighting the need for an efficient background reduction step with Cas9-mediated targeting. More recently, computational target enrichment strategies known as adaptive sequencing have been introduced through ReadFish ^15^ and UNCALLED ^16^, where non-target reads are “rejected”, allowing only target sequences to pass through the pores and be sequenced fully. However, these approaches required the DNA to be sheared to 8-15kb, impacting the ability to detect larger variants.

To address these limitations, we made modifications to the nCATS protocol to facilitate longer target capture and sequencing by reducing background fragments. We developed an <u>A</u>ffinity-based <u>C</u>as9-<u>M</u>ediated <u>E</u>nrichment method (ACME) that uses HisTag Isolation and Pulldown to remove background DNA. nCATS uses High-Fidelity (HiFi) Cas9 Nuclease that contains a 6-Histidine tag at its C-terminal. After the Cas9 cleavage and dA tailing step in nCATS, the Cas9 enzyme remains bound to the PAM-distal end i.e., non-target DNA side. Cas9 binding affinity is stronger at the PAM-distal end than the PAM-proximal side of the cleavage site ^17^, thereby protecting the non-target DNA end from subsequent sequencing adapter ligation. By introducing His-Tag specific magnetic beads at this step, we were able to pull down Cas-9 bound non-target fragments from the sample, allowing more target DNA to make it onto the flowcells **(Fig. 1)**. We designed a size titration panel consisting of important cancer genes of different size ranges – 10, 20, 40, 80, 150kb – to identify the largest possible target that could be effectively captured and sequenced as contiguous single fragments using ACME. We used ACME on this panel in two breast cell lines – MCF 10A ^18^ and SK-BR-3 ^19^ and compared its performance with nCATS for Cas9 targeting in MCF 10A and with PacBio and ONT whole genome sequencing in SK-BR-3 to detect SVs.

**Fig 1.**
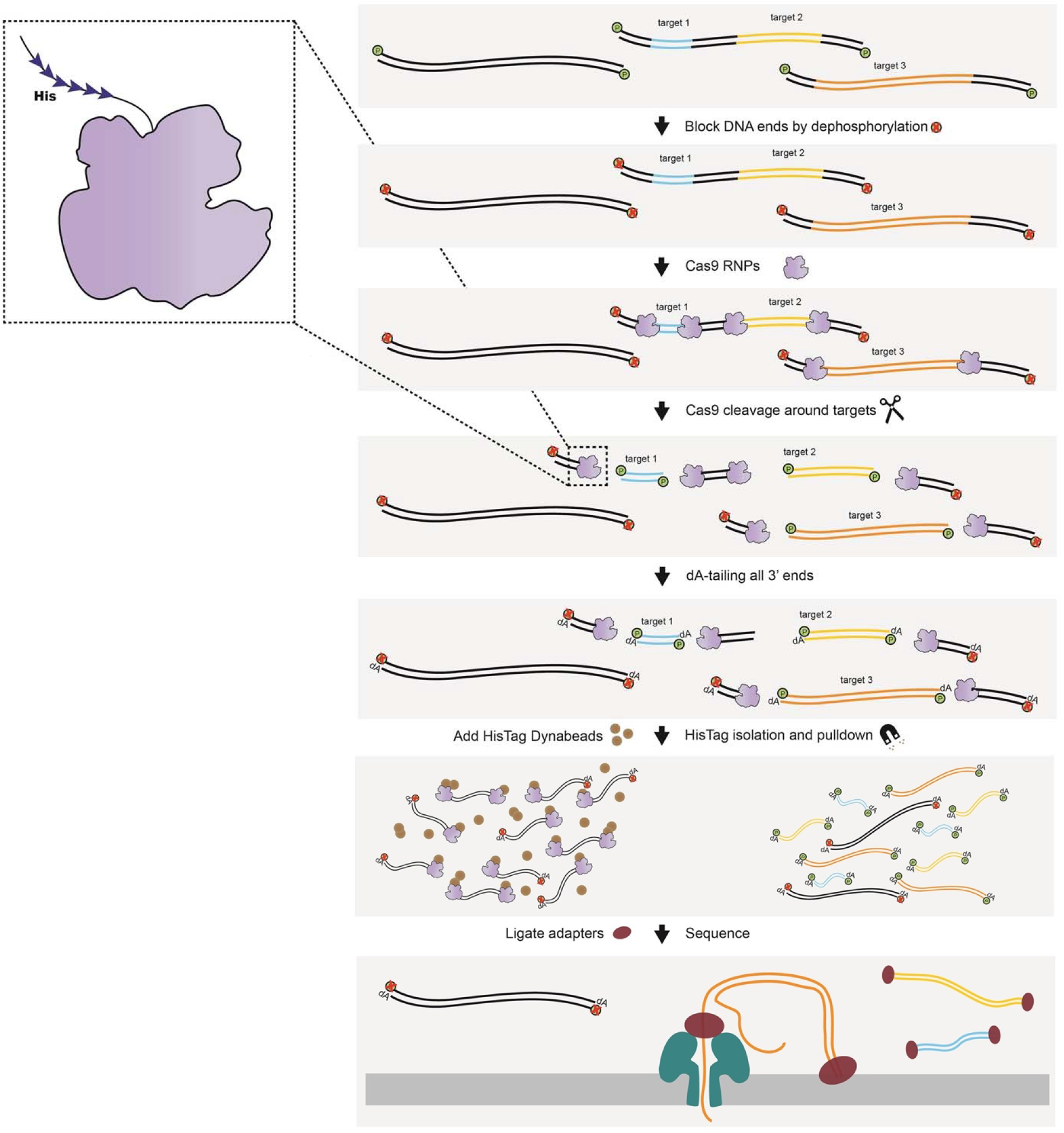
Affinity-based Cas9-Mediated Enrichment (ACME) Workflow. ACME aliows for the targeting of multiple regions of interest within the same reaction with single target read lengths as large as 1Oūkb. For ACME, individual samples starting with 5μg HMW DNA are prepped as follows:

- *Dephosphorylation:* 5’ ends of DNA are dephosphorylated to prevent the ligation of sequencing adapters to non-target strands.
- *Cas9 binding and cleavage:* Target specific crRNAs are bound with tracrRNA to form Cas9 RNPs. These RNPs cleave at either end of the targets, revealing blunt ends with now ligatable 5’ phosphates. After cleavage, the Cas9 enzyme stays bound to the PAM distal end on the non-target flanks, protecting those 5’ ends from adapter ligation.
- *dA-tailing:* All 3’ ends of DNA are dA-tailed to prepare the blunt ends for sequencing adapter ligation.
- *HisTag isolation and pulldown:* The Cas9 enzyme used for cleavage has a 6-Histidine tag at its C-terminal end (inset). Using His-Tag Dynabeads™ we isolate and pulldown Cas9 and the non-target DNA bound to it, thereby reducing background non-target reads.
- *Adapter ligation and sequencing:* Adapters are ligated to either side of the targets and the resultant Cas9 enriched library is loaded onto the flowcell for sequencing.

## Results

### Targeting a 200kb region around BRCA1

For our first pass of nCATS we used guides designed by Sage Sciences that targeted a 200kb region around the *BRCA1* gene ^20^ in SK-BR-3 to test the upper limits of potential target sizes using this method. This run generated a single 198kb read spanning the entire BRCA1 region and somewhat higher on-target coverage **(Fig. 2A).** This library was made from a single sample prep starting with 5μg of HMW DNA. We also replicated these efforts in MCF 10A **(Table 1, Supplemental Fig. 1A)**, a normal breast cell line. To improve on-target reads **(Table 1)**, we modified nCATS to incorporate higher DNA inputs and to reduce background/non-target reads.

**Fig. 2.**
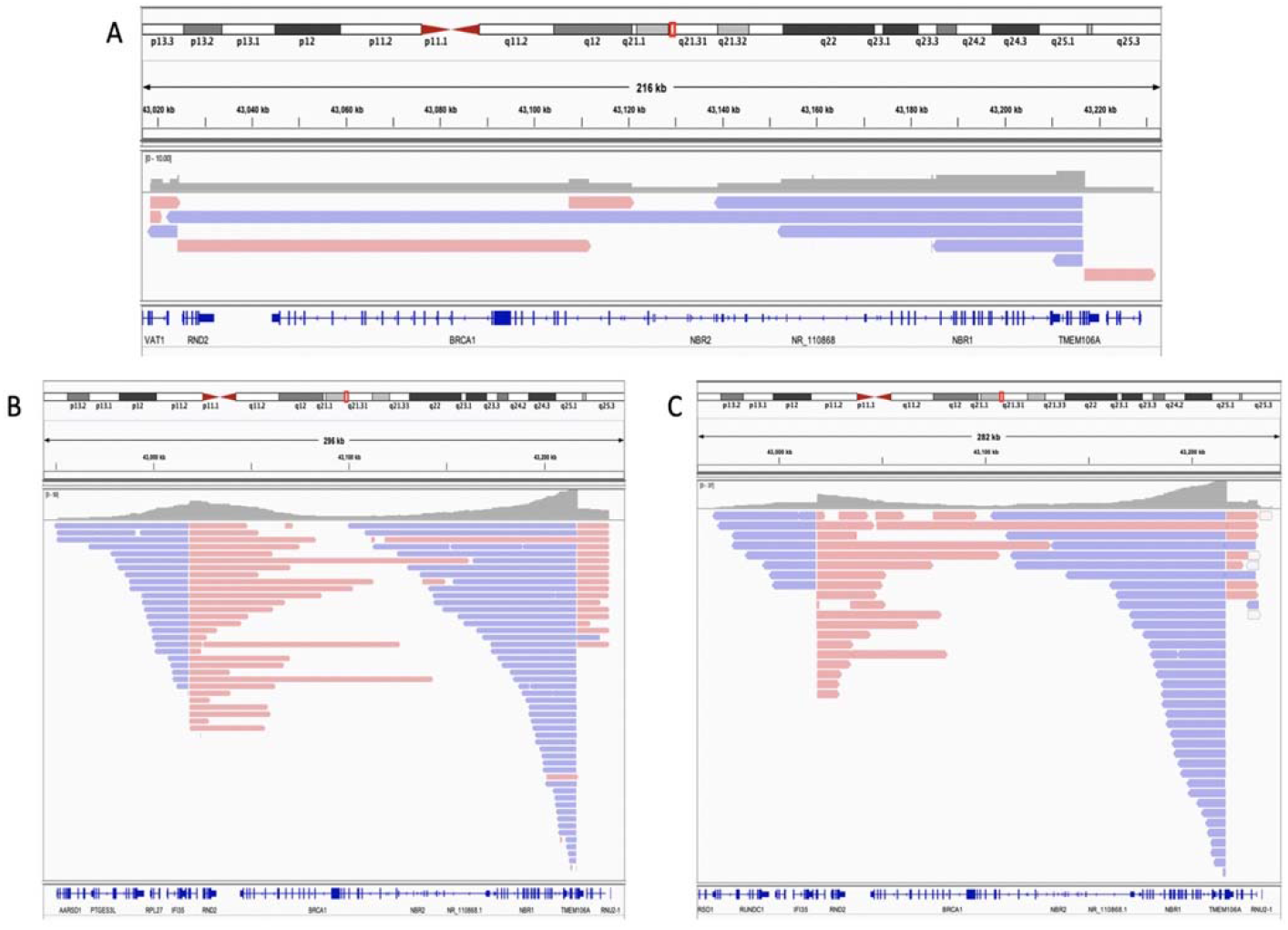
IGV plots representing reads that mapped to the *BRCA1* gene in different Cas9-mediated targeting libraries prepared from MCF 10A & SK-BR-3 DNA. Plus strand reads are shown in pink and minus strand reads in blue. **A:** Single sample library prep using SK-BR-3 DNA (nCATS). **B:** Pooled library prepared using 4 identical preps of MCF 10A DNA with the incorporation of Circulomics Short Read Eliminator kit (SRE). **C:** Pooled library prepared using 4 identical preps of SK-BR-3 DNA with the incorporation of ACME.

**Table 1.**
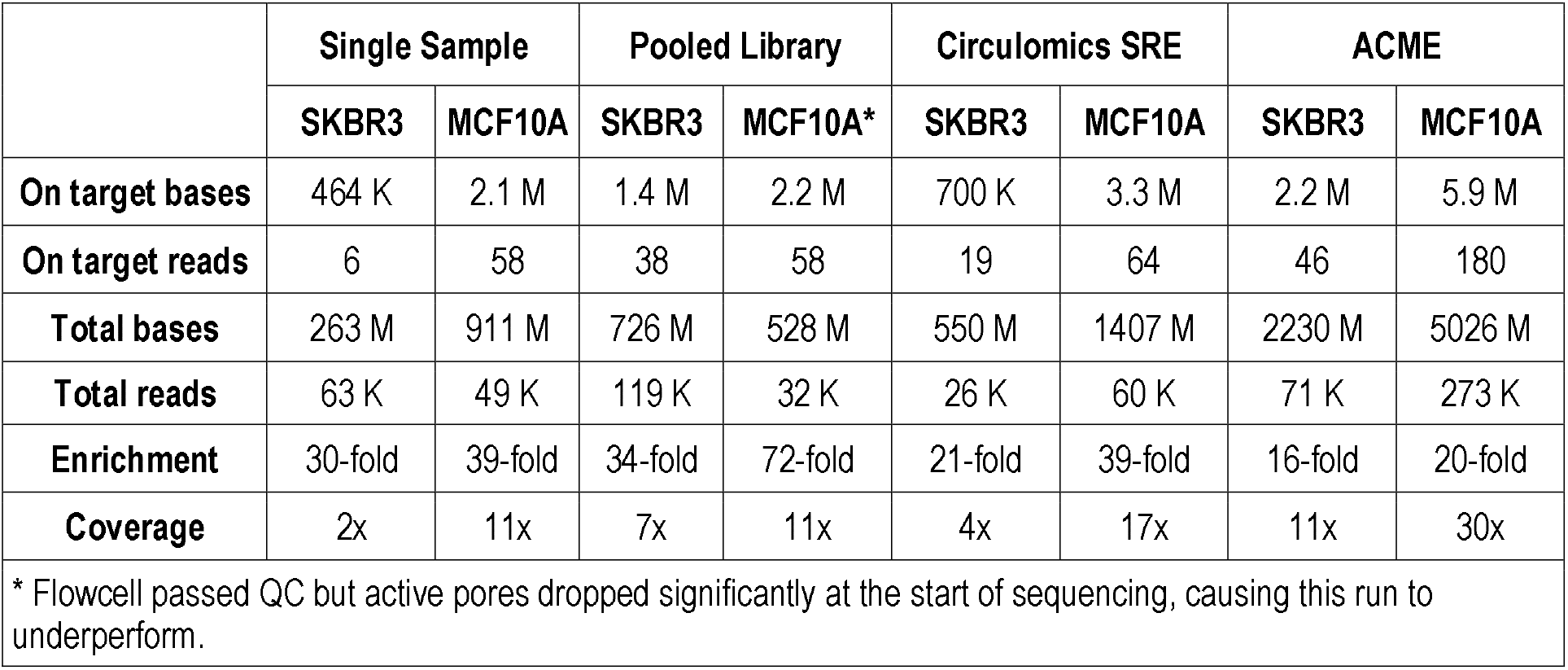
Run summaries for the different *BRCA1* Cas9-mediated targeted sequencing runs. Enrichment is relative to expected genome abundance for the total observed yield.

|  | Single Sample |  | Pooled Library |  | Circulomics SRE |  | ACME |  |
| --- | --- | --- | --- | --- | --- | --- | --- | --- |
|  | SKBR3 | MCF10A | SKBR3 | MCF10A* | SKBR3 | MCF10A | SKBR3 | MCF10A |
| On target bases | 464 K | 2.1 M | 1.4 M | 2.2 M | 700 K | 3.3 M | 2.2 M | 5.9 M |
| On target reads | 6 | 58 | 38 | 58 | 19 | 64 | 46 | 180 |
| Total bases | 263 M | 911 M | 726 M | 528 M | 550 M | 1407 M | 2230 M | 5026 M |
| Total reads | 63 K | 49 K | 119 K | 32 K | 26 K | 60 K | 71 K | 273 K |
| Enrichment | 30-fold | 39-fold | 34-fold | 72-fold | 21-fold | 39-fold | 16-fold | 20-fold |
| Coverage | 2x | 11x | 7x | 11x | 4x | 17x | 11x | 30x |

\* Flowcell passed QC but active pores dropped significantly at the start of sequencing, causing this run to underperform.

We tried three different modifications to improve depth across large targets of interest as described below:

*Pooled Library Preparation:* Given that we were targeting a single large region and nCATS involves no target amplification, four identical sample preps were pooled together to ensure sufficient copies of our target made their way to the sequencing pool. This prep gave us better target enrichment with 35-60 reads on target and a 35-70-fold enrichment **(Table 1)**. However, the on-target reads were not long enough to include the entire target region, with drops in coverage around the center of the region of interest **(Supplemental Fig. 1B)**.
*Circulomics Short Read Elimination (SRE):* We used the Short Read Eliminator kit from Circulomics Inc. to reduce potentially competing background reads <25kb in length and boost the coverage of our longer *BRCA1* fragments. While this modification did not work well for SK-BR-3 with only 19 reads on target **(Table 1, Supplemental Fig. 1C)**, the MCF 10A performed much better with one 142kb read and 64 reads on target **(Table 1, Fig. 2B)**.
*Affinity-based Cas9-Mediated Enrichment (ACME):* The Alt-R^®^ S.p. HiFi Cas9 Nuclease from Integrated DNA Technologies (IDT^®^) used in the ONT Cas9 protocol contains a C-terminal 6-His tag. Additionally, the Cas9 remains bound to the PAM-distal side (non-target DNA) after cleavage, protecting it from sequencing adapter ligation. By using Invitrogen™ Dynabeads™ His-Tag Isolation and Pulldown, we selected against the background by enriching for the non-Cas9 bound regions of interest **(Fig. 1)**. This modification gave us the most on target reads (50-180 reads) for *BRCA1* (198kb target) in both cell lines **(Table 1)**. While the MCF 10A run did not give us target spanning long reads **(Supplemental Fig. 1D)**, the SK-BR-3 ACME run gave us a 167kb read **(Fig. 2C)**.

The number of reads on target, irrespective of the nCATS modification used, were not sufficient to accurately call SVs in our region of interest. Based on these results, it seemed likely that a 200kb target size may have been too large a region to enrich for using this method, as evidenced by the clear drops in reads around the middle of the target region.

### Cancer Gene Panel

To identify the largest optimal region size that could be entirely spanned with single contiguous reads, we designed a size titration experiment to target a panel of genes belonging to different size ranges **(Table 2)**. These genes are present in the Catalogue Of Somatic Mutations In Cancer (COSMIC) database ^21,22^ and have known SVs in the HER-2-amplified breast cell line SK-BR-3 that were previously identified through whole genome long- and shortread sequencing ^19,23,24^. Target sizes ranged from 10kb to 150kb with a total targeted region of ~550kb **(Supplemental Table 1)**.

**Table 2.** COSMIC database genes with known SVs in SK-BR-3.

| Gene | Location | Gene size | Target size |
| --- | --- | --- | --- |
| MYC | 8q24.21 | 7517 | 12565 |
| HOXA9 | 7p15.2 | 12745 | 18506 |
| FGFR4 | 5q35.2 | 11270 | 19916 |
| STK11 | 19p13.3 | 22636 | 30286 |
| CDKN2A | 9p21.3 | 27291 | 30774 |
| TERT | 5p15.33 | 41880 | 44787 |
| KRAS | 12p12.1 | 46214 | 50955 |
| BRCA2 | 13q13.1 | 84192 | 91218 |
| PAX7 | 1p36.13 | 117860 | 120934 |
| APC | 5q22.2 | 138734 | 144265 |
| Total target size |  |  | 564206 |

Target size represents the shortest distance between crRNAs targeting the same gene

Given the improvement in the number of on-target bases and enrichment through pooling as well as background reduction, we further evaluated the performance of these modifications in targeting our cancer panel. Though iterative, Circulomics SRE has defined size cutoffs, making it a less amenable approach for background reduction when targets, such as our cancer panel, fall across different size groups. Since ACME relies on Cas9 bound nontarget DNA for effective background reduction, including more targets and therefore more crRNA guides per reaction results in more cut sites and better pulldown of background fragments.

As seen in **Supplemental Table 2**, all targets show a significant improvement in fold enrichment and coverage with the incorporation of ACME. **Fig. 3** shows reads that mapped to some representative genes of the panel in both MCF 10A and SK-BR-3 as viewed using IGV ^25^. Smaller genes on the panel like *MYC* (12kb, **Fig. 3A**) and *HOXA9* (18kb, **Fig. 3B**), and mid-sized genes like *STK11* (30kb, **Fig. 3C**) had decent depth even without ACME, but their coverage almost doubles with the incorporation of ACME. Larger gene targets such as *KRAS* (50kb, **Fig. 3D**) and *BRCA2* (90kb, **Fig. 4**), had poor coverage using nCATS. On using ACME, they not only have better coverage but also have significantly more target spanning reads that capture these genes in their entirety. Additionally, we see a drop in coverage around the middle of the target region for the two largest genes in the panel – *PAX7* (120kb, **Fig. 3E**) and *APC* (145kb, **Fig. 3F**), indicating an upper limit of 100kb for efficient targeting using ACME. Looking specifically at *BRCA2* **(Table 3)**, an important breast cancer gene, we see 60-90-fold enrichment of this 91kb target across both cell lines, giving us a coverage of 35-55x of the gene.

**Fig. 3.**
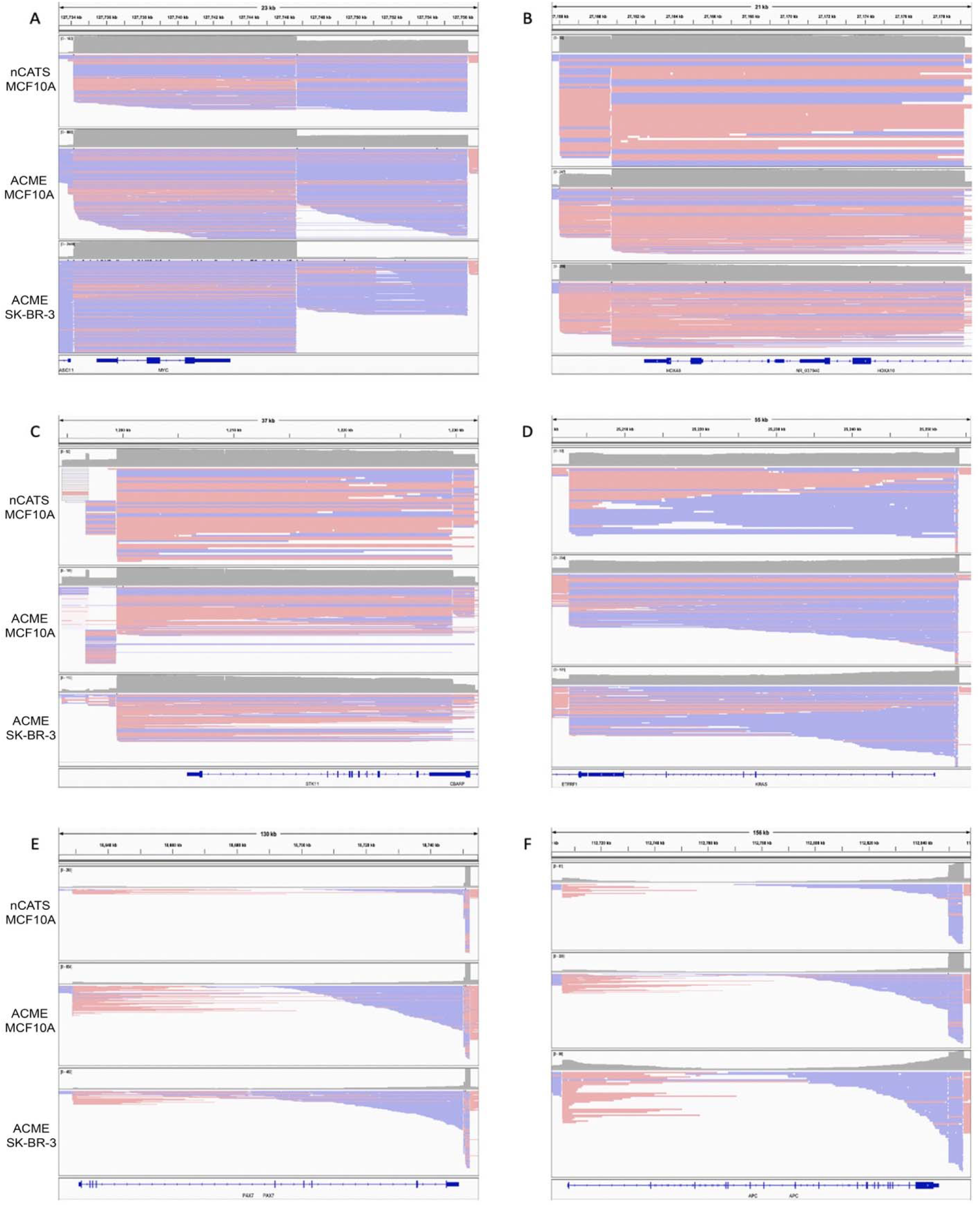
IGV plots showing a representative gene from each size range across both cell lines - MCF 10A and SK-BR-3. Plus strand reads are shown in pink and minus strand reads in blue. These genes were part of the 10 gene panel. For each image, the top panel shows reads that mapped to the gene in an MCF 10A library prepped without the ACME step (i.e., nCATS only); middle panel shows MCF 10A library prepped with the ACME step; bottom panel shows SK-BR-3 library prepped with ACME. Genes shown are **A**: *MYC* ~10kb **B**: *HOXA9* ~20kb **C**: *STK11* ~30kb **D**: *KRAS* ~50kb **E**: *PAX7* ~120kb and **F**: *APC* ~150 kb

**Fig 4.**
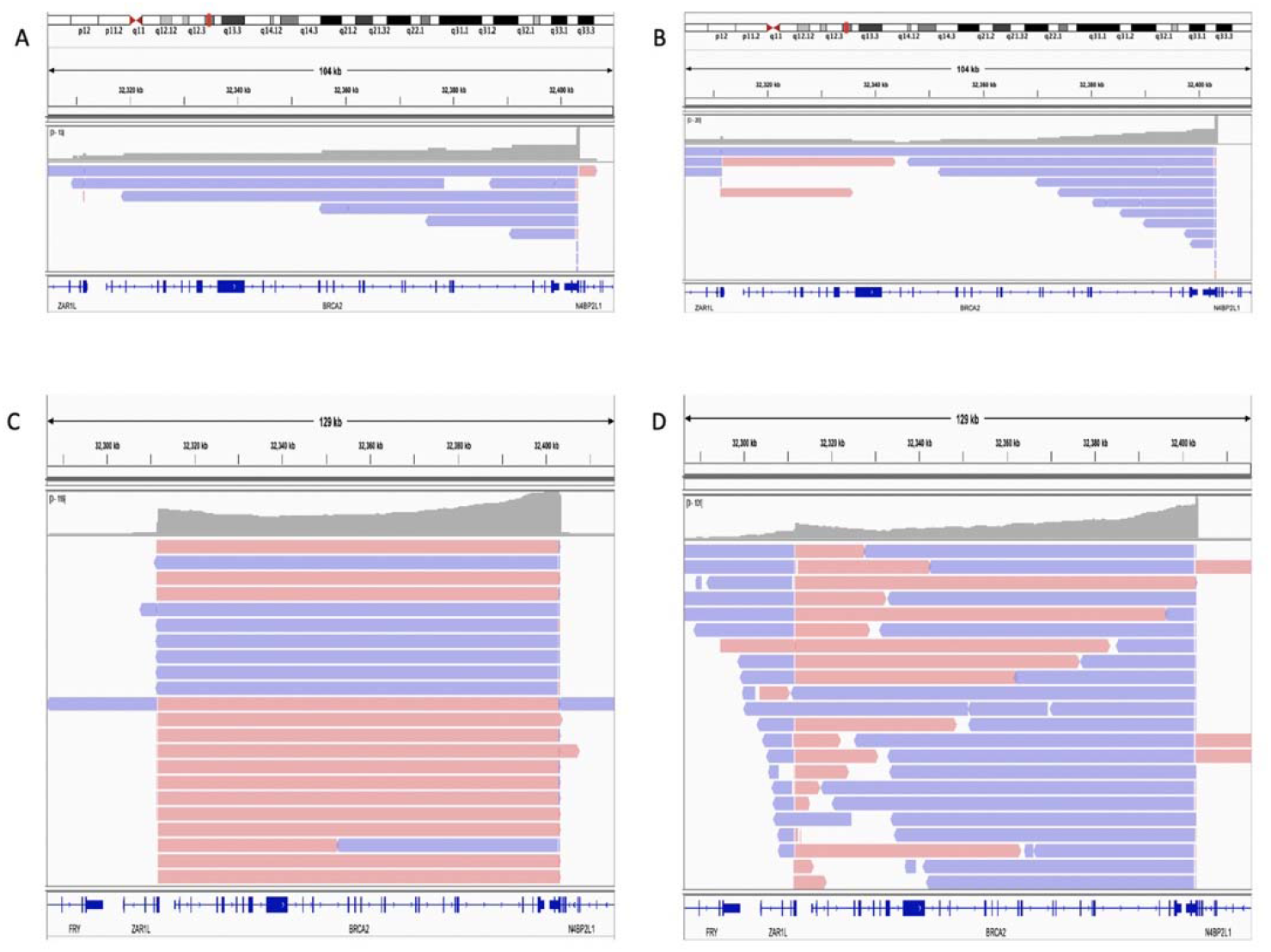
IGV plots showing reads that mapped to the *BRCA2* gene in different Cas9-mediated targeting libraries. Plus strand reads are shown in pink and minus strand reads in blue. **A**: Single sample library prep using MCF 10A DNA, without the ACME step (nCATS only) **B**: Library prepared by pooling together 3 identical library preps of MCF 10A DNA without the ACME step (nCATS only). **C-D**: Pooled library prepared from 4 identical preps of MCF 10A and SK-BR-3 DNA resp. with the incorporation of the ACME step to remove non-target DNA.

**Table 3.** Run summaries for *BRCA2* from the different targeted cancer panel sequencing runs.

| BRCA2 | Single MCF10A | Pooled MCF10A | ACME (Pooled) MCF10A | ACME (Pooled) SKBR3 |
| --- | --- | --- | --- | --- |
| On target bases | 326143 | 299552 | 5052621 | 3226173 |
| On target reads | 9 | 12 | 135 | 115 |
| Total bases | 190609890 | 276025707 | 2409003747 | 1894903771 |
| Total reads | 10299 | 17281 | 111274 | 102727 |
| Fold Enrichment | 101 | 41 | 86 | 62 |
| Coverage | 4x | 3x | 55x | 35x |
| For modifications where more than one run was performed during the testing phase, average values are represented in the table. |  |  |  |  |

One major advantage that ACME offers over other long-read targeting approaches is the ability to get multiple contiguous reads that span the entire length of the region of interest. For all genes on our panel that were <=100kb in size, we were able to obtain 3-7 times as many complete target spanning reads in comparison to the same regions captured using nCATS (**Table 4**). Even for larger targets, like *BRCA2* and *PAX7*, we obtained 2-20 reads spanning the entire 90-120kb targets, whereas these counts were between 0 to 2 reads for the ~100kb targets without ACME.

**Table 4.**
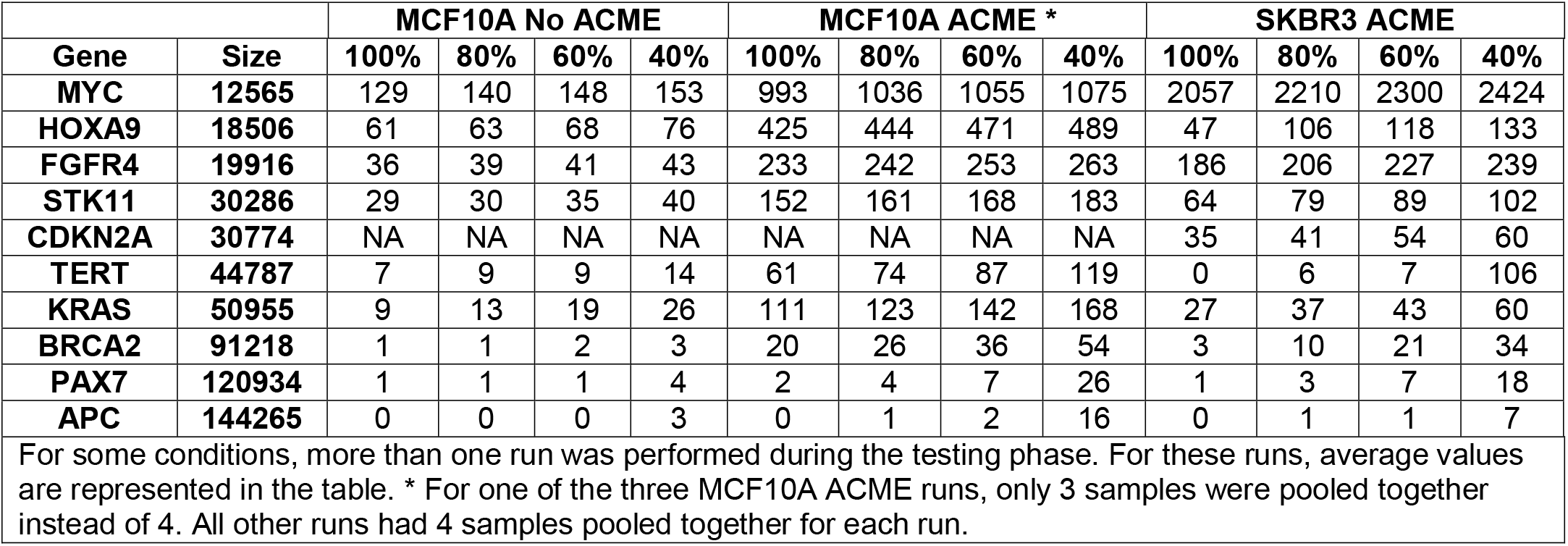
Number of reads spanning X% of gene for each targeted cancer panel gene across non-ACME and ACME pooled library sequencing runs.

We also prepared ACME enriched single libraries using just 5μg each of MCF 10A and SK-BR-3 DNA. On comparing these results with a single sample nCATS run without ACME (**Supplemental Table 3**), we see a significant increase in coverage using ACME, especially for the larger genes on the panel. As seen with the pooled runs, ACME also helps increase the number of single contiguous reads spanning the entire target (**Supplemental Table 4**). This is evident from **Supplemental Fig. 2**, which shows reads that mapped to a few representative genes from each size range on the panel. We clearly see that without ACME we lose the ability to capture reads that span the full length of the target in genes larger than 30kb, thereby limiting our scope of targeting these genes in rare samples.

### Structural variants

Having identified ACME as an effective targeted long-read method, we evaluated its performance in detecting SVs in SK-BR-3. While MCF 10A is a normal breast cell line ^18^, SK-BR-3 is a widely studied HER2+ breast cancer cell line ^19,24^. Furthermore, we have previously carried out whole genome sequencing of this cell line using short-read paired-end Illumina, long-read PacBio sequencing, 10X Genomics Linked Reads, and Oxford Nanopore sequencing ^23,26^ allowing us to compare ACME’s ability to detect variants with that of these widely used whole genome platforms. For ACME, SVs were called using Sniffles ^7^ and merged both manually and by using SURVIVOR ^27^, with the maximum distance between SVs set to 1000 bp and a minimum SV size of 20 bp, leaving us with 17 ACME SV calls. For the whole genome platforms, Sniffles and PBSV (https://github.com/PacificBiosciences/pbsv) were used to call SVs from ONT and PacBio datasets, while SVaBA ^28^, Lumpy ^29^, Manta ^30^, GROC-SVs ^31^, NAIBR ^32^, and LongRanger ^33^ were used on the Illumina/10X data to call SVs ^23,26^. We subsetted SVs inferred from whole genome data across each sequencing platform to include only those that appeared within our target coordinates for each of the ten genes on our panel, giving us 11 ONT Svs and 15 PacBio SVs. For Illumina, SVs supported by at least 2 of 6 callers used by Aganezov et al. were included, accounting for Illumina’s lower accuracy in SV detection and to reduce the false positives that are known to arise on inferring SVs from short-read data.

Overlaps between ACME and each whole genome SV callset were inferred based on their positions, such that an SV from the ACME callset that occurred within 1000 bp of an SV from a whole genome callset overlapped and were considered the same event. The same approach was used to determine overlaps between the different whole genome SV callsets. We observed very strong concordance between ACME SVs with those from whole genome ONT and PacBio **(Fig. 5)**. ACME successfully identified all SVs from the PacBio and ONT datasets that occurred within our target regions **(Supplemental Table 5)**. Additionally, ACME inferred 2 SVs that were not called by PacBio, and 6 SVs that were not called by ONT, which could be attributed to the higher on target coverage achieved by ACME. Of the 4 Illumina SVs, only 2 events overlapped with each ACME, PacBio, and ONT SV callsets, and were common between all platforms. Our results highlight that ACME performs similar to ONT and PacBio whole genome sequencing for comprehensive SV inference thus establishing it as a viable and effective targeted long-read approach. We also looked at ACME’s ability to detect SVs from a single sample prep with initial DNA input of 5 μg. ACME was able to identify 12 SVs from the single library data compared to the 17 SV calls from the pooled library, with 9 overlaps with ONT SV calls and 12 overlaps with PB SV calls. **(Supplemental Fig. 3)**

**Fig. 5.**
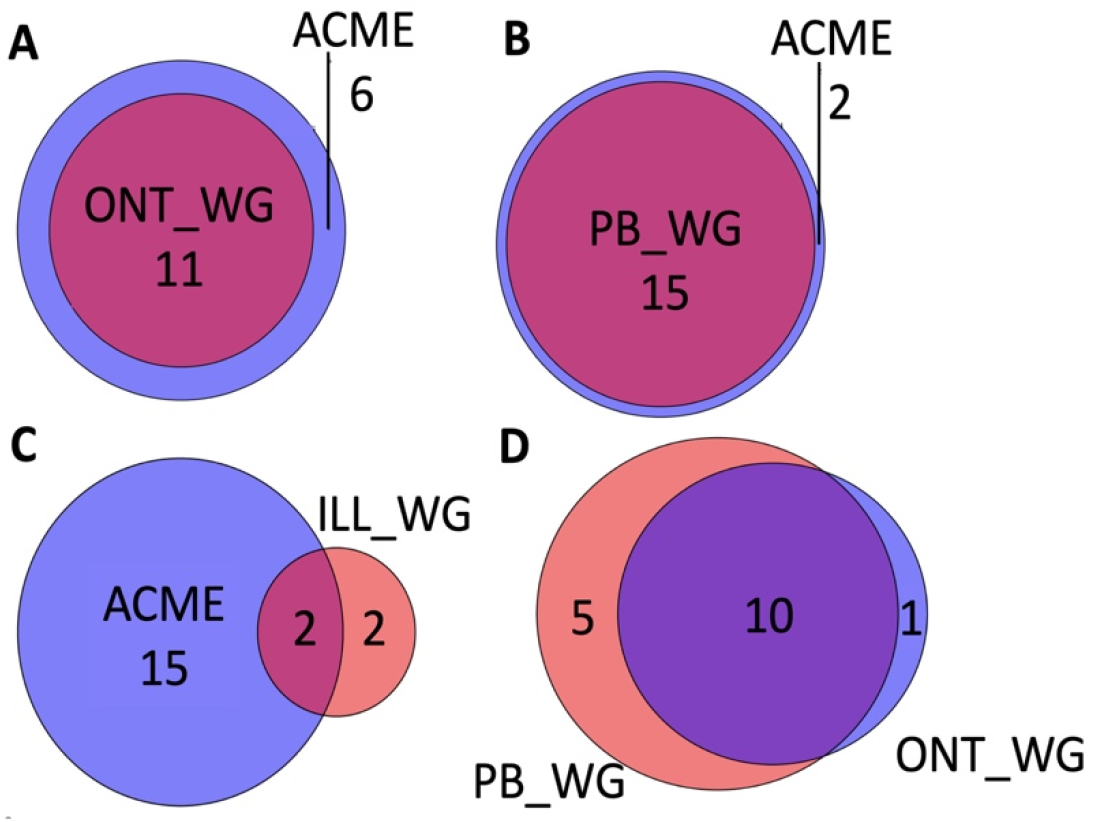
Comparison of pooled ACME with whole genome platforms for SV detection in SK-BR-3. **AC**: SVs detected by ACME that overlap with ONT, Pacific Biosciences (PB), and Illumina (ILL) whole genome sequencing (WG) resp. **D**: High concordance observed between PB and ONT WG for SV detection in SK-BR-3. **Note**: Only SVs appearing within our target coordinates for each of the ten genes on our panel were included in these comparisons. SVs were detected from pooled sample prep ACME run.

### Single Nucleotide Polymorphisms

Nanopore sequencing tends to have low sensitivity in SNP detection due to its difficulty in distinguishing signal events in repetitive regions like homopolymers. We explored how the increased coverage using ACME would improve the ability to call SNPs from the SK-BR-3 dataset. We used SNP calls from the short-read paired-end Illumina dataset, and also re-ran SNP callers on our previously sequenced long-read PacBio and Oxford Nanopore datasets ^23,26^ to look for variants common between the platforms.

All SNP callsets were filtered to include only those variants that fell within our target regions. xAtlas ^34^ was used to call SNPs from the Illumina data giving us 500 in-target SNPs. Clair ^35^ was used to call SNPs from the whole genome ONT (1234 SNPs) and PacBio (601 SNPs) datasets and the targeted ACME dataset (592 SNPs).

For the first round of comparisons between platforms, we looked at all SNPs that met a minimum depth of >= 10 for each platform. Assuming Illumina as the gold standard for SNP detection, we looked for SNPs from each long-read platform that overlapped with SNPs called from the Illumina dataset. As seen in **Fig. 6** ACME performed on par with the whole genome long-read platforms when compared to Illumina for SNP detection. We also noted a high number of ONT whole genome SNPs that did not overlap with Illumina (**Fig. 6A**). This could be explained by the overall lower depth of the ONT WG data and the older basecaller used on this dataset, which produced lower single read accuracy and could therefore generate more false positives due to basecall errors. In contrast, we observed that the higher depth from targeted ACME (**Fig. 6B**), called with the latest basecaller, helped bring down the number of false positive SNPs called by the nanopore platform. On performing the same set of comparisons using data generated from a single library prep of ACME (5 μg initial DNA input), we observed that of the 337 SNPs inferred by ACME, 151 overlap with Illumina. **(Supplemental Fig. 4)**.

**Fig. 6.**
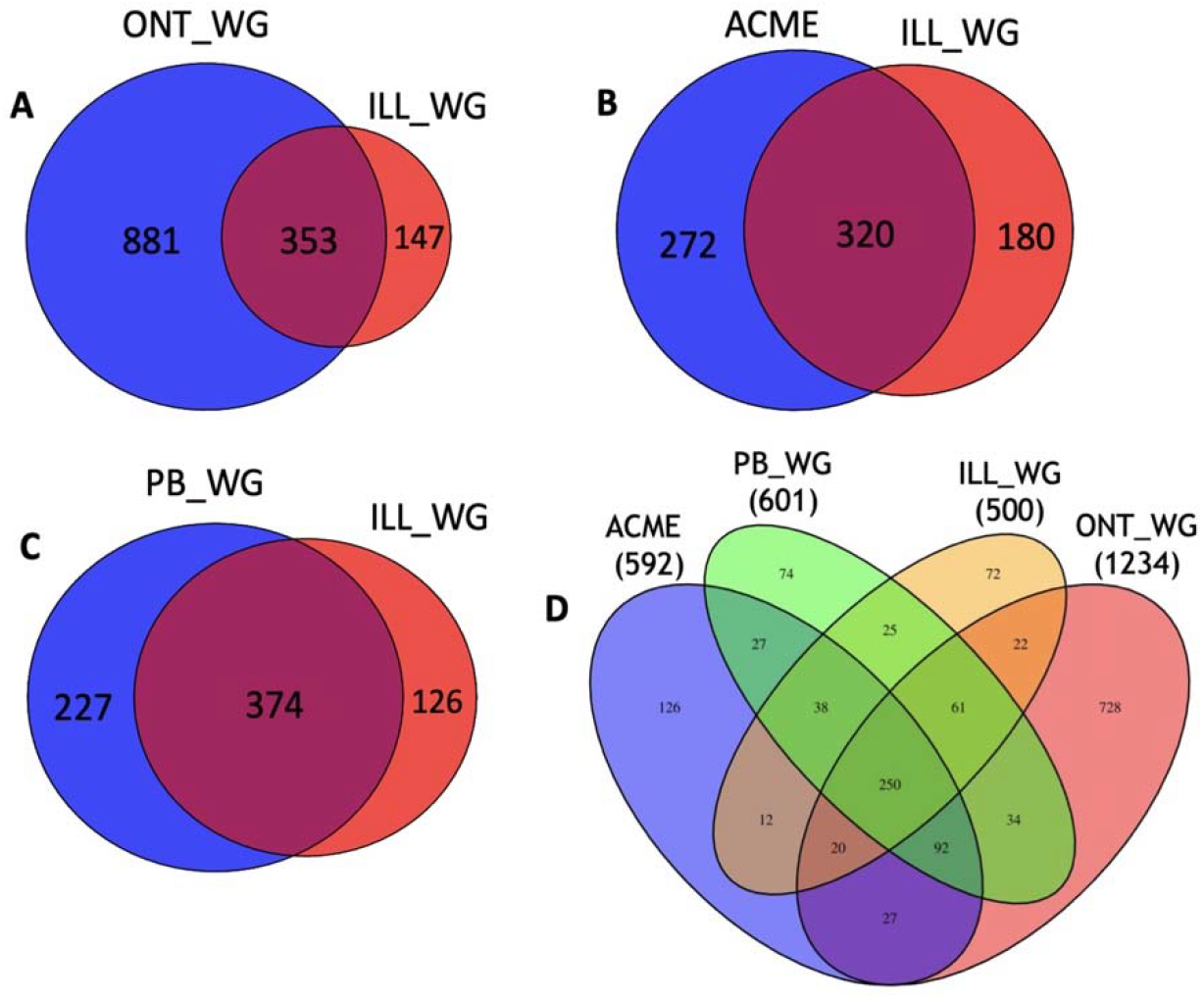
Comparison of long-read platforms with Illumina whole-genome sequencing for SNP detection in SK-BR-3. **A-C**: SNPs detected by Illumina (ILL) that overlap with ONT, ACME, and Pacific Biosciences (PB) resp. **D**: SNPs detected by and that overlap between each platform. **Note**: Only SNPs with minimum depth of 10x appearing within our target coordinates for each of the ten genes on our panel were included in these comparisons. SNPs were detected from pooled sample prep ACME run.

Given the lack of coverage across the full length of targets >100kb in the ACME dataset, we reran the above comparisons between platforms (minimum depth >=10) after filtering out SNPs present in the *PAX7* (~120kb) and *APC* (~150kb) genes across all platforms. We observed an increase in the percentage of ACME-ILL overlapping SNPs from 64% to 76%. The overlaps between ONT-ILL and PB-ILL however remained consistent (**Supplemental Fig. 5**). This indicates that ACME is as capable as, if not better than, the whole genome long-read platforms for SNP detection in regions with complete coverage of target without any dropouts in the middle.

Lastly, in addition to calling SNPs *de novo* as described thus far, we also looked at forced genotype calls using the Illumina SNP set as the query. A forced genotype approach is particularly useful for detecting specific SNPs of interest in long-read datasets. Through the forced genotype approach, we specifically looked for 618 SNPs called from the Illumina dataset (minimum depth >=6) in our long-read datasets to see if these platforms detected the query SNPs irrespective of their depth. We found that ACME detected 88% of SNPs in the query set, again performing on par with PacBio whole genome (90% Illumina SNPs detected) and ONT whole genome (87% Illumina SNPs detected). Additionally, we observed that forced genotype calls generated from the single library ACME run were also largely comparable, with 82% of Illumina SNPs detected **(Supplemental Fig. 6**).

## Discussion

Whole genome, long-read sequencing is currently expensive and too low throughput for large-scale variant studies. Targeted DNA sequencing is a cost-effective approach that facilitates higher sample throughput and improves accuracy, giving depths necessary to detect rare alleles in a heterogenous population of cells. In this study, we developed an <u>A</u>ffinity-based <u>C</u>as9-<u>M</u>ediated <u>E</u>nrichment (ACME) approach, that builds upon the existing nCATS method by adding a simple background reduction step, physically increasing the on-target to off-target ratio in the sample prior to sequencing. Unlike exome capture, ACME helps capture entire gene regions, including introns, as single sequencing reads up to 100kb in size. ACME leverages the presence of a 6-histidine tag at the C-terminal domain of the Cas9 nuclease that is used for cleaving and releasing the targets from the dephosphorylated DNA. Since Cas9 remains preferentially bound to the non-target side of the cleavage site, using HisTag isolation and pulldown we were able to physically remove Cas9-bound non-targets from the sequencing pool prior to adapter ligation. Using ACME, we captured and sequenced fragments as large as 100kb with 35-55x coverage, doubling the existing single read sizes capturable. By eliminating smaller competing non-targets from the sequencing library, we saw a 3-to 7-fold increase the number of whole gene spanning reads per target, thereby aiding in SV discovery by reducing potential mapping errors.

Compared to traditional nCATS, we find that ACME not only helps in increasing target coverage but also allows for the complete spanning of genes by generating reads as long as 100kb - up to 3 times the targetable size and read lengths achieved using nCATS. We have shown that even for 90-120kb sized targets (*BRCA2* and *PAX7*), ACME helped capture 2-20 reads spanning the entire targets, whereas without ACME these counts were between 0 to 2 reads. We ran the nCATS and ACME experiments parallelly using the same HMW DNA, since DNA quality is important for optimal targeting and generating long reads. Our results therefore specifically highlight the advantage offered by ACME’s ability to reduce competing background reads. Two other Cas9-based targeting approaches that involve background reduction - CaBagE ^13^ and Negative Enrichment ^14^ - use exonuclease digestion to eliminate non-target DNA. Both studies have only successfully achieved target read lengths of <35kb in size and pose the additional risk of digesting nicked target fragments that are not protected by Cas9 binding. Furthermore, the exonuclease step adds 2-4 hours to the library prep, while the ACME step adds <45 mins, making it a quick and efficient targeted long-read approach.

We demonstrated ACME’s ability to detect variants by comparing it with PacBio, ONT, and Illumina whole-genome sequencing of SK-BR-3. For SVs, we showed that ACME detected all SVs within our targets that were previously inferred by PacBio and ONT whole-genome sequencing ^23,26^. ACME’s higher coverage also helped identify a few novel SVs previously undetected by whole genome PacBio and ONT, which we are currently validating. Since the limitations for SV detection with long-read technologies are largely cost driven, we show that ACME performs on par with not only the more expensive PacBio platform, but also surpasses depths achievable by ONT whole-genome sequencing. For SNPs, we found that ACME performs similar to whole genome PacBio and ONT when we compared the long-read platforms to Illumina, which is considered the gold standard. To perform a fairer comparison between platforms, we excluded SNPs from *PAX7* and *APC* across all platforms on finding coverage dropouts in targets >100kb in the ACME callsets. We then found that ACME not only matches but surpasses the long-read platforms in their ability to detect SNPs. ACME detected more SNPs called by Illumina than the whole-genome PB and ONT, likely due to increased target coverage. We also found that ACME performs on par with these platforms when we adopted the forced genotype approach. These results are promising as they show that ACME can effectively replace the whole genome long-read platforms for SNP detection by offering better depth and increased sensitivity at a fraction of the cost.

While pooled ACME runs start with 20 μg of initial DNA input, we also evaluated the performance of single library ACME preps starting with 5 μg to account for the realistic possibility of having limited sample DNA available. For SV detection, we observed that ACME still performs better than ONT whole-genome sequencing, while performing on par with PacBio by failing to detect only 3 SVs found by PacBio. For SNPs, the number of ACME-Illumina overlaps fell drastically on switching from pooled to single ACME preps. This drop in variant calls, especially SNPs, may be attributed to the lower coverage observed in the single library data. While all experiments described in this study were run using the early access developer protocol (ONT protocol versions ENR_9084_v109_revB_04Dec2018 and ENR_9084_v109_revH_04Dec2018) that required sourcing individual reagents separately, with the release of the standalone ONT Cas9 kit (SQK-CS9109), coverage from single library preps is no longer a concern. We tested the CS9109 kit performance with and without ACME across different preps and concluded that this newer kit offers improved yield and a much higher coverage for the single as well as pooled library preps than previously observed. Our efforts to run ACME on the low pore Flongle device have been unsuccessful and ONT has withdrawn support for nCATS on the Flongle.

Adaptive sequencing approaches like ReadFish ^15^ and UNCALLED ^16^ can target 100s of regions per run, however increased rejection of reads affects sequencing yield, leads to quicker pore burnouts, and lower target coverage (<60x, usually close to 20-30x) than enzymatic methods can achieve. Since sheared DNA (8-15kb) is required for these approaches, it is better suited to detect single-nucleotide variants (SNVs), copy number variants (CNVs), and repeat expansions than detecting larger rare variants ^36^. On evaluating if ACME could be combined with adaptive sequencing to play to both their strengths, we found that adaptive sequencing offers no enrichment, and in fact negatively impacts enzymatic enrichment, for panels with few targets (<0.1-1% of the genome). Therefore, while adaptive sequencing may not be the best tool for discovery of rare or large variants, it is great for clinical settings ^36^ to survey 100s of targets for known variants, which can then be confirmed with higher depth enzymatic approaches such as nCATS or ACME.

A current limitation of ACME is the inability to sequence deeply targets >100kb in size without coverage dropouts in the center. However, this could be a function of DNA size more than the targeting approach itself and may be solved by switching to the recently released Ultra HMW (UHMW) DNA Extraction protocol from Circulomics (Pacific Biosciences). Moreover, even with the 100kb size limit ACME offers distinct advantages over the other Cas9 based targeting approaches. Targets >100kb can be captured using ACME with the designing of tiled guides, with tiles spanning 50-80kb regions as opposed to 20-30kb with the other approaches. This will not only reduce the number of tiles per target, but also the crRNA guides per reaction, allowing for more targets to be captured per reaction. The main advantage ACME offers over other amplification free long-read targeting approaches is the ability to capture several contiguous reads, up to 100kb in size, that span the whole target from start to end. This helps in reducing mapping errors and aid in SV detection even with lower target coverage for large gene targets.

## Methods

### Cell culture and DNA extraction

MCF 10A and SK-BR-3 cell lines were obtained from ATCC and cultured as per standard guidelines. High Molecular Weight (HMW) DNA was extracted using an adaptation of a modified Sambrook and Russell ^37^ DNA extraction described by Jain *et al*. ^38^ Approximately 3×10^7^ cells were spun at 150 rpm for 10 min to pellet. The cells were resuspended in 200 μl PBS. 10 ml TLB was added (100 mM NaCl, 10 mM Tris-Cl pH 8.0, 25 mM EDTA pH 8.0, 0.5% (w/v) SDS, 20 μg/ml Qiagen RNase A), vortexed at full speed for 5 s and incubated at 37°C for 1 hr. 100 μl Proteinase K (Qiagen, final conc. 200 μg/ml) was added and mixed by slowly rotating end over end three times followed by 2 hr. incubation at 50°C with gentle mixing every 30 mins. The lysate was split into two, phenol-purified using 5 ml phenol-chloroform-isoamyl alcohol (25:24:1), mixed end over end for 10 min, and spun at 4000 rpm for 10 mins. Supernatant was transferred to fresh tubes using wide bore tips, 5 ml chloroform-isoamyl alcohol (24:1) was added, mixed end over end for 10 min, and spun at 4000 rpm for 10 mins. Using wide bore tips, supernatants were combined and DNA was precipitated by the addition of 4 ml 5 M ammonium acetate and 30 ml ice-cold 100% ethanol followed by gentle rocking for 15 min. DNA was recovered with a glass hook followed by washing twice in 70% ethanol. After spinning down at 10,000g, ethanol was removed followed by air drying for 15 min. 300 μl of nuclease free water was added to the DNA and left at 4°C overnight to resuspend. DNA was quantified using the Qubit fluorometer (ThermoFisher Scientific) and Femto Pulse System (Agilent) before use.

### Guide RNA design

crRNA probes were designed to target and cut on either side of the region of interest (ROI) to excise that region. Four to six crRNA guides were designed per target for redundancy – 2-3 upstream of the ROI targeting the (+) strand and 2-3 downstream that targeted the (-) strand. Guides were designed using CHOPCHOP (http://chopchop.cbu.uib.no/) ^39,40^ and chosen based on efficiency (>0.3), GC content (40-80%), self-complementarity (score=0), and predicted mismatches (<5), with at least 1kb of flanking sequence either side of the ROI. Duplex format of guide RNAs were ordered, comprising of synthetic crRNAs (IDT Cas9 Alt-R™ crRNAs, custom designed) and tracrRNAs (IDT Alt-R™, Cat #1072532). Guide sequences for all genes in the panel are provided in **Supplementary Table 1**.

### RNP assembly and Cas9 cleavage

All crRNAs for all genes in the panel were resuspended at 100 μM in TE (pH 7.5) and pooled into an equimolar mix prior to guide RNA assembly. As described in Gilpatrick *et al*. ^41^ and using ONT protocol versions ENR_9084_v109_revB_04Dec2018 and

ENR_9084_v109_revH_04Dec2018 for reference, crRNA mix and tracrRNA were combined such that the tracrRNA and total crRNA concentration were both 10 μM. gRNA duplexes were formed by denaturation at 95°C for 5 min and then cooling to room temp for 5 min. Ribonucleoprotein complexes (RNPs) were constructed by combining 10μl of annealed crRNA•tracrRNA pool (10 μM) with 0.8μl of HiFi Cas9 Nuclease V3 (62 μM, IDT Cat #1081060) in 10 μl of 10X CutSmart Buffer (NEB, Cat #B7204) at a final volume of 100 μL, incubated for 30 min at room temperature, then stored on ice until use. This mix is good for 10 reactions and assembled RNPs can be stored at 4°C for up to a week. Genomic DNA was dephosphorylated by adding 3 μl of Quick CIP enzyme (NEB, Cat #M0508) to 5 μg of input DNA (in total volume of 24 μl at >210 ng/μl) with 3 μl 10X CutSmart Buffer (NEB, Cat #B7204), and incubating for 10 min at 37°C, followed by heating for 2 min at 80°C. After allowing the sample to return to room temp, 10 μL of the pre-assembled Cas9 RNP was added to the sample. In the same tube, 1μL of 10mM dATP (NEB, Cat # N0440S) and 1uL of Taq DNA polymerase (NEB, Cat #M0273) were added for A-tailing of DNA ends. The sample was then incubated at 37°C for 15 min for Cas9 cleavage followed by 5 min at 72°C for A-tailing. For pooled libraries, four identical preps of 5μg each were prepared parallelly and taken to the next step. For single libraries, only 1 prep of 5 μg was carried forward to the next step.

### nCATS modifications for BRCA1 targeting

#### • Pooled Library Preparation

For each sequencing run, four reactions were set up starting with 5μg HMW DNA for each reaction, giving a total of 20μg starting material across the 4 reactions. Each reaction was taken though the steps on nCATS individually until the adapter ligation step, wherein each product was eluted to a lower volume and pooled together at the end before loading onto the flowcell. In order to avoid overloading the flowcell, only half the library was loaded initially with the other half loaded 24 hours into the sequencing run.

#### • Circulomics Short Read Elimination (SRE)

Since the pooled library prep generated more on target reads, we started with four identical preps each of MCF 10A and SK-BR-3 DNA. For each prep, SRE was performed immediately after the Cas9 cleavage and dA-tailing step. The products of SRE were cleaned up using 1X Ampure XP beads, eluted in a smaller volume, and pooled together respectively for both MCF 10A and SK-BR-3.

#### • Affinity-based Cas9-Mediated Enrichment (ACME)

For both cell lines, four identical sample preps of 5μg HMW DNA each, were run through our modified nCATs protocol described in detail below. Like the SRE modification, products of the Cas9 cleavage and dA-tailing step were immediately processed using ACME to remove Cas9 bound non-target DNA from the sample. ACME products were cleaned up using 1X Ampure XP beads, eluted in a smaller volume, and pooled together respectively for both MCF 10A and SK-BR-3.

### Affinity-based Cas9-Mediated Enrichment (ACME)

For pooled ACME preps, four identical preps of 5μg each were set up for both MCF 10A and SK-BR-3. crRNA guides for all 10 genes were pooled together into an equimolar mix for the Cas9 cleavage step. ACME was performed on each sample separately as follows: The 42μl cleaved sample from the Cas9 cleavage and dA-tailing step was resuspended in 700μl of freshly prepared 1X binding/wash buffer (2X Binding/Wash Buffer: 100 mM Sodium Phosphate pH 8.0, 600 mM NaCl, 0.02% Tween™-20). Invitrogen^™^ His-Tag Dynabeads^™^ (Cat #10103D) were thoroughly resuspended in the vial (vortexed >30 sec), 50 μL (2 mg) of which was transferred to a fresh 1.5μl Eppendorf DNA LoBind tube. Tube was placed on a magnet for 2 min, supernatant was discarded, and the sample (prepared in 1X binding/wash buffer) was added to the beads and mixed well. After incubating on a roller for 5-10 min at room temperature, the tube was placed on a magnetic rack for 2 min, and the supernatant was carefully transferred to a fresh tube using wide bore tips. *Note: The targets/regions of interest are suspended in the supernatant while the Cas9-bound non-targets are bound to the His-Tag beads. DO NOT discard the supernatant at this step*. The sample was cleaned up using 1X Ampure XP beads (Beckman Coulter, Cat #A63881) and eluted in nuclease-free water. Based on the number of initial parallel preps, the elution volume was adjusted to give a final volume of 44μl at the end of ACME i.e., eluted in 11 μl each (for the pooled library) or 44 μl (for single library). ACME products were quantified using the Qubit fluorometer (ThermoFisher Scientific). 42μl of the ACME enriched sample was taken forward to the adapter ligation step.

### Library prep and sequencing

Sequencing adaptors and ligation buffer from the Oxford Nanopore Ligation Sequencing Kit (ONT, LSK109) were ligated to cut DNA ends using T4 DNA ligase (NEBNext Quick Ligation Module E6056) for 10 min at room temperature. The sample was cleaned up using 0.3X Ampure XP beads (Beckman Coulter, Cat #A63881) and washed twice with the long-fragment buffer (LFB; ONT, LSK109), resuspending the beads in the buffer each time. Sample was eluted in 15 μl of elution buffer (EB; ONT, LSK109) for 30 min at room temperature. *Note: Elution time of 30 min is recommended for targets >30kb*. Sequencing libraries were prepared by adding 25 μL sequencing buffer (SQB; ONT, LSK109) and 13 μl loading beads (LB; ONT, LSK109) to 12 μl of the eluate. Each sample was run on a FLO-MIN106 R9.4.1 flow cell with >1200 active pores following Platform QC on the GridION sequencer. 30 μl of flush tether (FLT; ONT, LSK109) was added to 1 tube of flush buffer (FB; ONT, LSK109). Initial flow cell priming was performed with 800 μL of this mix and allowed 5 min to equilibrate. The flow cells were then primed with 200 μL of the priming mix prior to loading the sequencing libraries.

### Analysis

Nanopore reads were base called using Guppy v3.2. Reads were transferred to an isilon NL400 storage server and mapped to the hg38 human reference genome (UCSC Genome Browser) using either MiniMap2 ^42^ or NGMLR ^7^. The BAM files were sorted and indexed with Samtools ^43^ and on-target reads were visualized using IGV ^25^. Capture metrics including depth and breadth of target coverage and target enrichment were generated with Picard HSmetrics (https://broadinstitute.github.io/picard/index.html) and GATK DepthofCoverage ^44^. Structural variants were called with Sniffles ^7^. VCF files were sorted with VCFtools ^45^. SNPs were called using Clair ^35^. Variant comparisons between datasets and target-spanning read counts were done with BEDtools ^46^.

#### • SV detection

Sniffles uses strand information to help confirm the event type as such: Strand orientation of the adjacency (DEL/INS:+-, DUP:-+, INV:++/−−). During merging using SURVIVOR, SV types were not required to be the same, but strand/orientation was required to match. For ACME, ONT and PacBio, we did iterative filtering based on read support, which was required to be >=4 reads supporting the SV, and variant read counts, which was required to be >= 5% of total read counts.

#### • SNP detection

##### SNP calling with xAtlas on Illumina data

The “PASS” filters were engaged during SNP calling, initially with a minimum depth >=6, variant read depth >2, and a strand bias filter, giving us a total of 618 SNPs within our targets. Depth was then raised to >=10, giving us 500 SNPs that were used for the comparisons. The following additional filters were used for SNP calling with xAtlas on the Illumina data:

“low_snpqual”, Description= “SNP logistic regression P-value is > 0.5”
“low_VariantReads”, Description= “Variant read depth is > 2”
“low_VariantRatio”, Description= “Variant read ratio is not less than Phred-scaled Genotype Likelihood scores cutoff”
“low_coverage”, Description= “Total coverage is >= 6”
“high_coverage”, Description= “Total coverage is <= 8000”
“single_strand”, Description= “All variant reads are NOT in a single strand direction”
“No_data”, Description= “At least 1 valid read on this site”
“No_var”, Description= “At least 1 valid variant read on this site”

##### SNP calling with Clair on the long-read data

Minimum depth was set to >= 10 with 10% alternate allele frequency and a filtration score of 400. Variants are divided into 10 categories: 1) a homozygous reference allele; 2) a homozygous 1 SNP allele; 3) a heterozygous 1 SNP allele, or heterozygous 2 SNP alleles; 4) a homozygous 1 insertion allele; 5) a heterozygous 1 insertion allele, or heterozygous 1 SNP and 1 insertion alleles; 6) heterozygous 2 insertion alleles; 7) a homozygous 1 deletion allele; 8) a heterozygous 1 deletion allele, or heterozygous 1 SNP and 1 deletion alleles; 9) heterozygous 2 deletion alleles; and 10) a heterozygous 1 insertion and 1 deletion alleles. The likelihood value of the 10 categories is calculated for each candidate variant, and the category with the largest likelihood value is chosen. The variant quality is calculated as the square of the Phred score of the distance between the largest and the second-largest likelihood values.

## Notes

### Competing Interest Statement

- SVI has received travel bursaries from Oxford Nanopore Technologies supporting oral and poster presentations at conferences.
- WRM is currently affiliated with the following organizations:
Cold Spring Harbor Laboratory, Orion Genomics, Gerson Lerman, State University of New York Stony Brook, New York Genome Center (NYGC)

