## Supplemental Figures for "ACME: an Affinity-based Cas9 Mediated Enrichment method for targeted nanopore sequencing"

Supplemental Figure 1

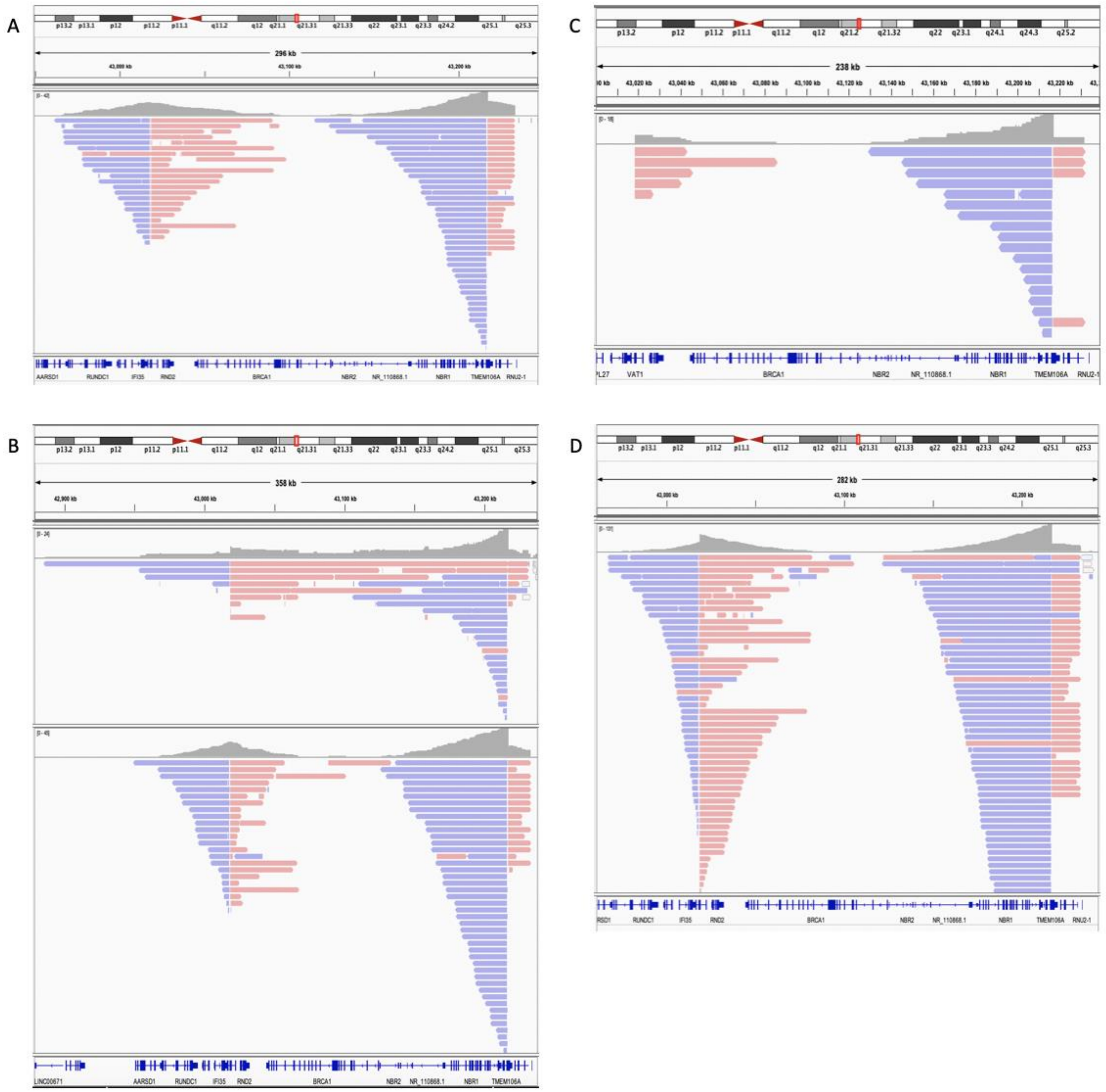


**Supp. Fig. 1** represents reads that mapped to the *BRCA1* gene in different Cas9-mediated targeting libraries prepared from MCF 10A & SK-BR-3 DNA. **A:** Single sample library prep using MCF 10A DNA. **B:** Libraries prepared by pooling together 3 identical library preps of SK-BR-3 (top) and MCF 10A (bottom) DNA. **C:** Pooled library prepared using 4 identical preps of SK-BR-3 DNA with the incorporation of Circulomics Short Read Eliminator kit (SRE). **D:** Pooled library prepared using 4 identical preps of MCF 10A DNA with the incorporation of ACME.

Supplemental Figure 2

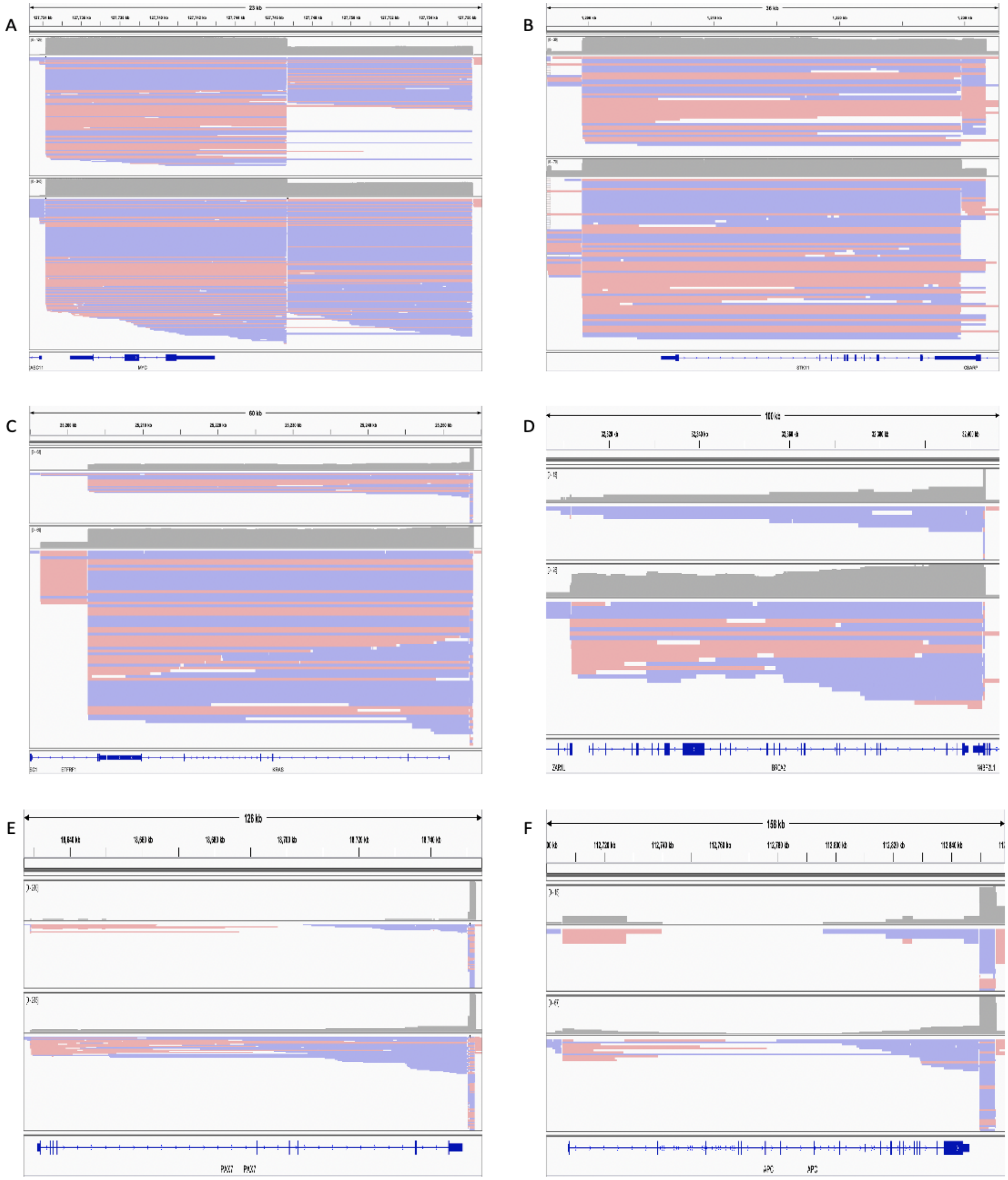

**Supp. Fig. 2** IGV plots showing representative genes from different size ranges in single library nCATS and ACME runs prepped using only 5µg MCF 10A DNA each. Plus strand reads are shown in pink and minus strand reads in blue. These genes were part of the 10 gene panel. For each image, the top panel shows reads that mapped to the gene in an MCF 10A library prepped without the ACME step (i.e., nCATS only) and bottom panel shows MCF 10A library prepped with the ACME step. Genes shown are **A**: *MYC* ~10kb **B**: *STK11* ~30kb **C**: *KRAS* ~50kb **D**: *BRCA2* ~90kb **E**: *PAX7* ~120kb and **F**: *APC* ~150 kb

Supplemental Figure 3

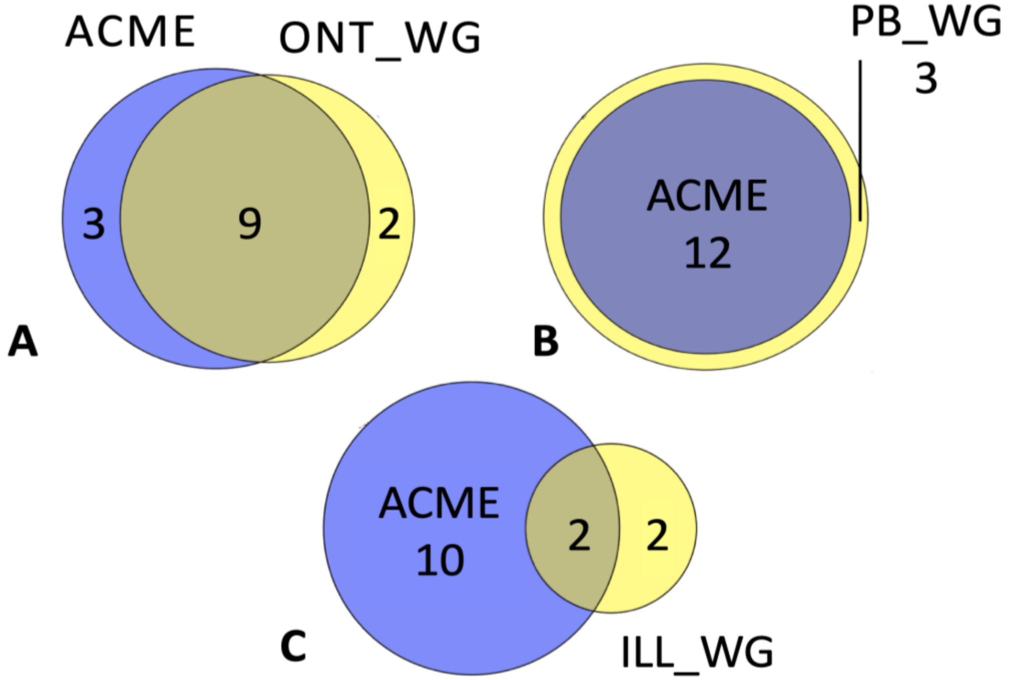


**Supp. Fig. 3. Comparison of single ACME with whole genome platforms for SV detection in SK-BR-3. A-C**: SVs detected by ACME that overlap with ONT, Pacific Biosciences (PB), and Illumina (ILL) whole genome sequencing (WG) resp. **Note**: Only SVs appearing within our target coordinates for each of the ten genes on our panel were included in these comparisons. SVs were detected from single sample prep ACME run.

Supplemental Figure 4

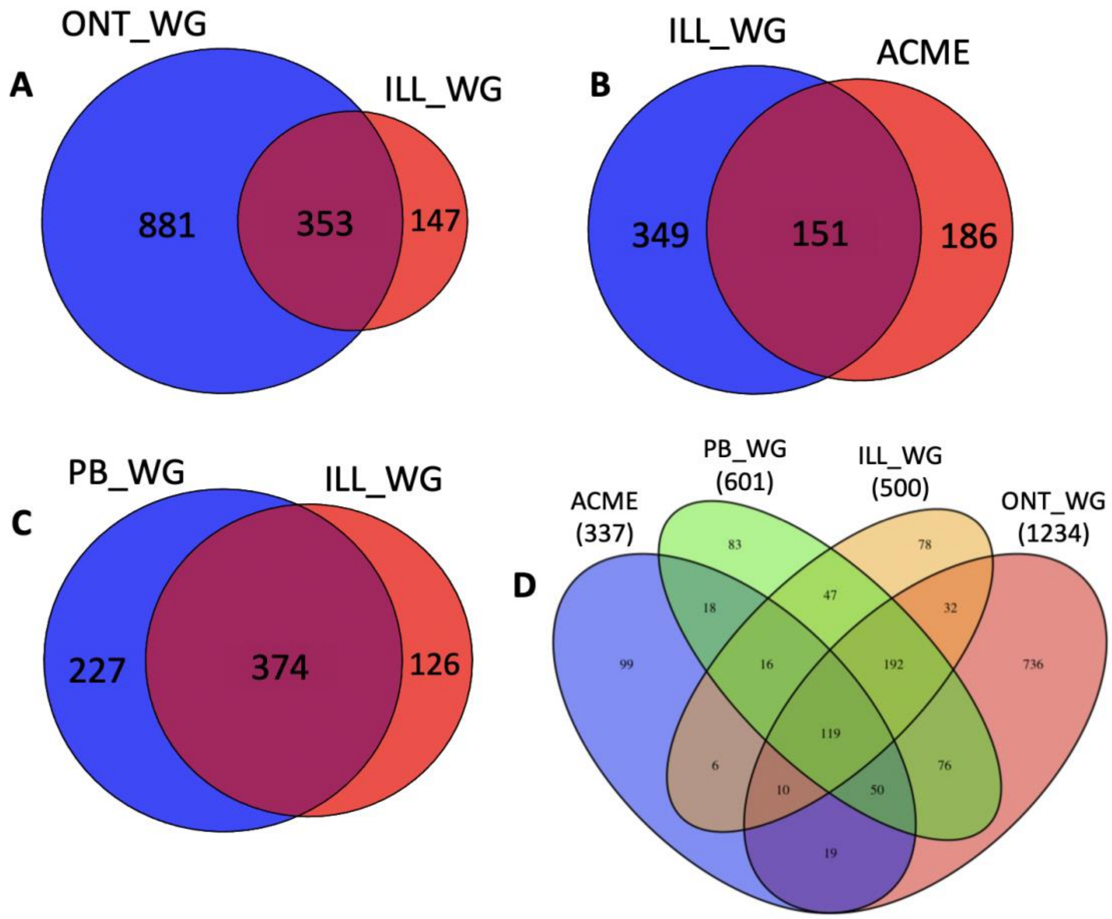


**Supp. Fig. 4. Comparison of long-read platforms with Illumina whole-genome sequencing for SNP detection in SK-BR-3. A-C**: SNPs detected by Illumina (ILL) that overlap with ONT, ACME, and Pacific Biosciences (PB) resp. **D**: SNPs detected by and that overlap between each platform. **Note**: Only SNPs with minimum depth of 10x appearing within our target coordinates for each of the ten genes on our panel were included in these comparisons. SNPs were detected from single sample prep ACME run.

Supplemental Figure 5

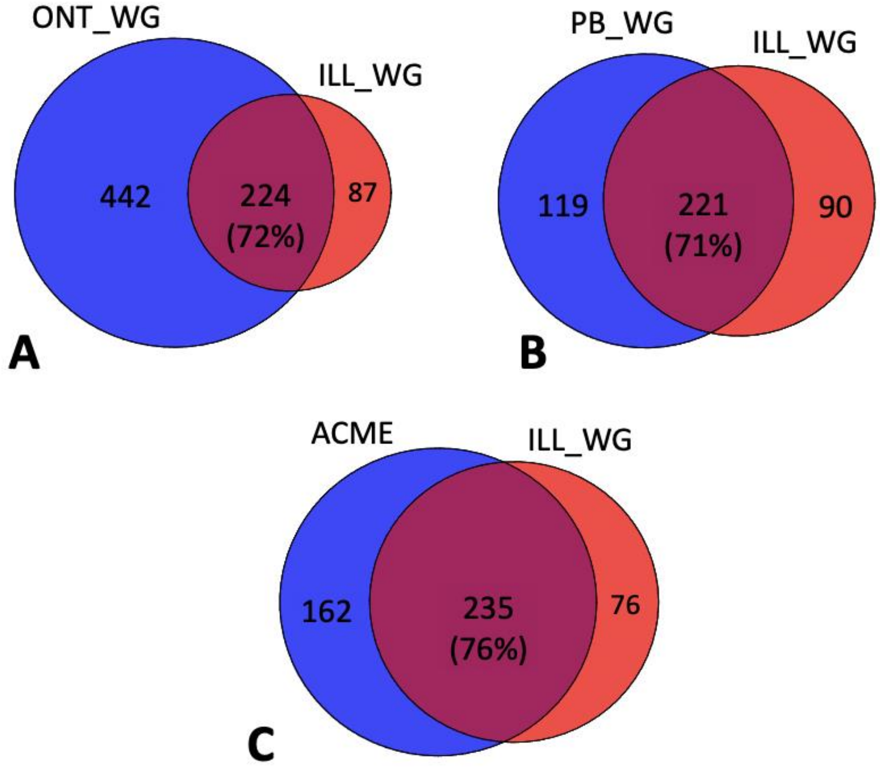


**Supp. Fig. 5. Comparison of long-read platforms with Illumina whole-genome sequencing for SNP detection in SK-BR-3 with exclusion of *PAX7* and *APC* SNPs. A-C**: SNPs detected by Illumina (ILL) that overlap with ONT, Pacific Biosciences (PB), and ACME resp. **Note**: Only SNPs with minimum depth of 10x appearing within all our targets except for *PAX7* and *APC* genes were included in these comparisons. SNPs were detected from pooled ACME run.

Supplemental Figure 6

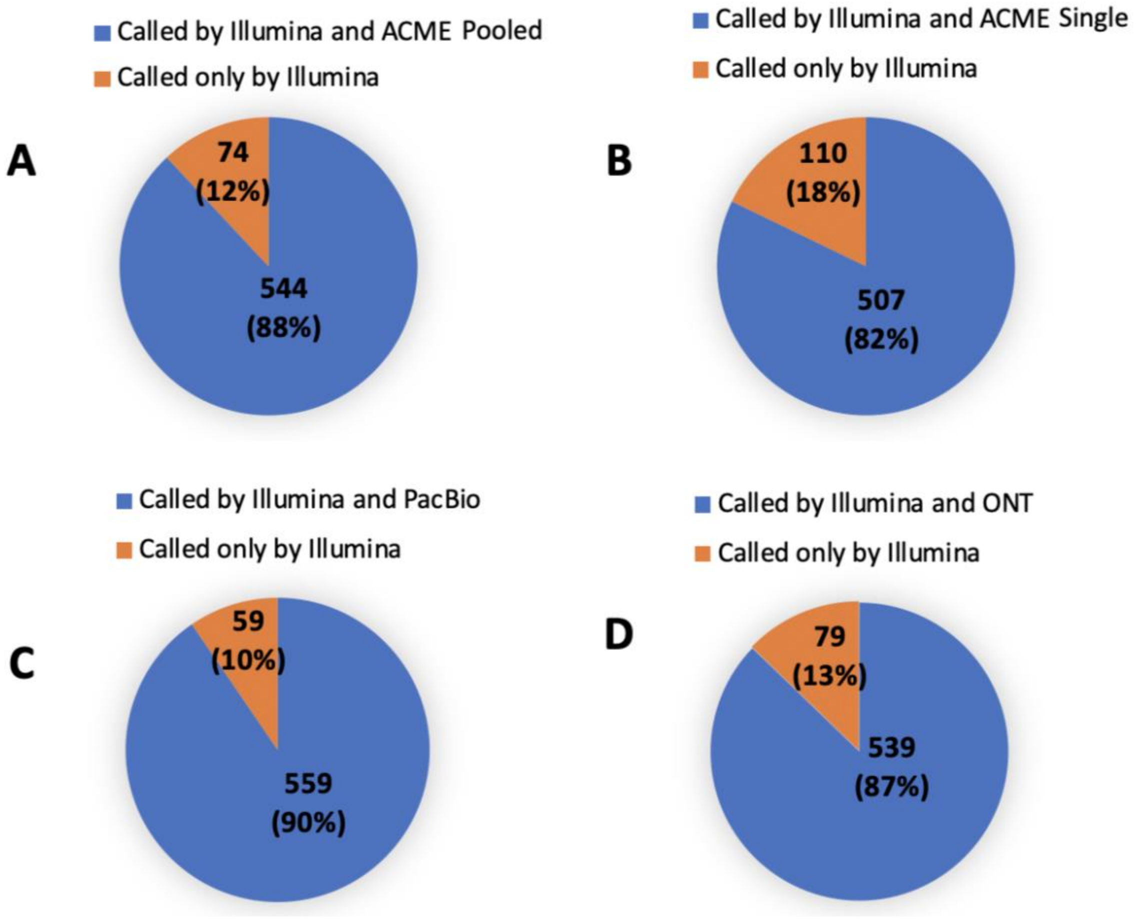


**Supp. Fig. 6. Forced genotype calls across long-read platforms using Illumina SNP set as query. A-B**: Illumina (ILL) SNPs detected in Pooled ACME and Single ACME targeted runs resp., irrespective of their depth in each run. **C-D**: Illumina (ILL) SNPs detected in Pacific Biosciences (PB) and ONT whole genome runs resp., irrespective of their depth in each run. **Note**: Only Illumina SNPs appearing within our targets that had a minimum depth of 6x were included in these comparisons.
