## Supplemental Tables 1-4 for "ACME: an Affinity-based Cas9 Mediated Enrichment method for targeted nanopore sequencing"

Supplemental Table 1

**Supplemental Table 1. crRNA guide coordinates (GRCh38/hg38) used to target cancer gene panel**

| Guide | Loci | Sequence (5’-3’) # | Strand | Guide | Loci | Sequence (5’-3’) # | Strand |
|---|---|---|---|---|---|---|---|
| MYC_L4 | chr8:127734131 | TACTTTCGCAAACCTGAACGCGG | + | MYC_R4 | chr8:127756321 | CTATCAGCTGTGTTGCGAGTGGG | - |
| MYC_L5 | chr8:127733817 | GCCATTACCGGTTCTCCATAGGG | + | MYC_R5 | chr8:127756368 | ATACTTCGAGACAGTAAGAGTGG | - |
| FGFR4_L6 | chr5:177079925 | TATCCGAGACCAGTGAATAAGGG | + | MYC_R6 | chr8:127746696 | AAACTTCATCAAGCGATCCAGGG | - |
| FGFR4_L7 | chr5:177079813 | CGTGCATGCCAACACCCGGTGGG | + | FGFR4_R7 | chr5:177106005 | AGTACCAGCGCGAGAACGTGAGG | - |
| HOXA9_L8 | chr7:27157986 | CCATAAACCCGACCTCACAATGG | + | FGFR4_R8 | chr5:177100991 | TGGAGCATCAGAGCTCGCATAGG | - |
| HOXA9_L9 | chr7:27160714 | TGGGCGGTCATCAAGTTCTGGGG | + | FGFR4_R9 | chr5:177099841 | GGGCTTTCGGGAGTTAGTGGGGG | - |
| STK11_L10 | chr19:1194527 | TCTTGCCCTGCGCTCCCGAAGGG | + | HOXA9_R10 | chr7:27185958 | CGGATCAGTGACAAACCGCGGGG | - |
| STK11_L11 | chr19:1199476 | AGATATCTACGAGTTCGGTGGGG | + | HOXA9_R11 | chr7:27179220 | TAAGGCGAGCTTATTCCCAGGGG | - |
| CDKN2A_L12 | chr9:21961369 | GGCACTCTACCAAATCACGAAGG | + | STK11_R12 | chr19:1229762 | TACGCCAGCATCGAGGGCCGCGG | - |
| CDKN2A_L13 | chr9:21965556 | TGTGCTCGGGAAACACGCGTGGG | + | STK11_R13 | chr19:1231739 | GTCCGCACCCACCTGACCCGGGG | - |
| TERT_L14 | chr5:1250590 | GCACCTGGCACCAACATCGACGG | + | CDKN2A_R14 | chr9:21997024 | GTTGTTAAGCAACCCCATGACGG | - |
| TERT_L15 | chr5:1251835 | TCCTCACAGCTCCTATACGGGGG | + | CDKN2A_R15 | chr9:21996330 | TGAGTGAAATCTACCTACCGGGG | - |
| KRAS_L16 | chr12:25196365 | ACTCAAGTTAGGATTCATCGGGG | + | TERT_R16 | chr5:1296622 | GATCTTCATTGAATGCCGGGAGG | - |
| KRAS_L17 | chr12:25202651 | AGTAGAGTGTGTGCGCCGAATGG | + | TERT_R17 | chr5:1297465 | GTGACCACCTGTTATCCCATGGG | - |
| BRCA2_L20 | chr13:32311322 | GCTTGCTCTTCGTCTCCTGAGGG | + | KRAS_R18 | chr12:25253606 | CGGCTTCTAAGAAAGTACCCAGG | - |
| BRCA2_L21 | chr13:32311721 | TAAGCTGCGCCCTCTTGTAAGGG | + | KRAS_R19 | chr12:25254110 | TACGTGCACACCTTTGTACGTGG | - |
| PAX7_L24 | chr1:18628891 | GCCATACCAAGGCAACCTGGAGG | + | BRCA2_R23 | chr13:32402939 | GTTGTCACCATGGATATTAGAGG | - |
| PAX7_L25 | chr1:18629339 | GATCTCCCAATCTCTCCCGGGGG | + | BRCA2_R24 | chr13:32403452 | TCTTGAAGCACGTAATTGAGAGG | - |
| APC_L26 | chr5:112705267 | GGGGGTCTAAGGTATCAGCTGGG | + | PAX7_R28 | chr1:18750273 | GACTTACCTGAAGGATAGATTGG | - |
| APC_L27 | chr5:112705521 | GCCCTTTGAGGTCCTACACAGGG | + | PAX7_R29 | chr1:18750818 | GCGTGCAGATCAATGAAACCAGG | - |
| | | | + | PAX7_R30 | chr1:18752350 | GGGAGGGTTAACTAAAACTATGG | - |
| | | | + | APC_R31 | chr5:112849786 | TTTTATGGCGATGGTGTCAGAGG | - |
| | | | + | APC_R32 | chr5:112855467 | GTACACCTCACGTTGAACCTTGG | - |

### Sequence includes NGG at 3’ end.

#### Supplemental Table 2

**Supplemental Table 2. Run summaries for the targeted cancer panel genes across non-ACME and ACME pooled library sequencing runs**

| | | Bases | | | Reads | | | Fold enrichment | | | Coverage | | | % Bases over 60x | | |
|---|---|---|---|---|---|---|---|---|---|---|---|---|---|---|---|---|
| **Gene** | **Size** | **MCF10A No ACME** | **MCF10A ACME *** | **SKBR3 ACME** | **MCF10A No ACME** | **MCF10A ACME *** | **SKBR3 ACME** | **MCF10A No ACME** | **MCF10A ACME *** | **SKBR3 ACME** | **MCF10A No ACME** | **MCF10A ACME *** | **SKBR3 ACME** | **MCF10A No ACME** | **MCF10A ACME *** | **SKBR3 ACME** |
| **MYC** | **12565** | 1771340 (1404259 – 2138421) | 12879331 (5325104 – 22769613) | 28571726 | 167 (125 – 209) | 1142 (465 – 2041) | 2656 | 1787 (1449 – 2125) | 1480 (785 – 2347) | 4012 | 141 (112 – 170) | 1025 (424 – 1812) | 2274 | 99.7 (99.6 – 99.8) | 100.0 (100 – 100) | 100 |
| **HOXA9** | **18506** | 737004 (632890 – 841118) | 4559611 (1477889 – 8042888) | 4121256 | 45 (42 – 48) | 248 (57 – 481) | 212 | 519 (387 – 650) | 340 (210 – 563) | 393 | 40 (34 – 45) | 246 (80 – 435) | 223 | 0.0 | 99.6 (98.7 – 100) | 100 |
| **FGFR4** | **19916** | 1372826 (976086 – 1769565) | 9308550 (3906162 – 16411419) | 2468116 | 93 (62 – 124) | 584 (251 – 1018) | 179 | 844 (757 – 932) | 677 (357 – 1067) | 219 | 69 (49 – 89) | 467 (196 – 824) | 124 | 49.7 (0 – 99.3) | 100.0 (99.9 – 100) | 99.7 |
| **STK11** | **30286** | 1095712 (844295 – 1347129) | 5307532 (1518349 – 9276905) | 2831087 | 52 (35 – 68) | 227 (79 – 390) | 139 | 454 (379 – 530) | 237 (155 – 397) | 165 | 36 (28 – 44) | 175 (50 – 306) | 93 | 0.0 | 72.2 (16.8 – 100) | 98.9 |
| **CDKN2A** | **30774** | 95 (0 – 190) | NA | 1679376 | 0 (0 – 0) | NA | 42 | 0 | NA | 96 | 0 (0 – 1) | NA | 55 | 0.0 | NA | 12.9 |
| **TERT** | **44787** | 567938 (542824 – 593051) | 4701177 (3941630 – 5193107) | 4039285 | 21 (19 – 22) | 121 (124 – 155) | 122 | 172 (113 – 230) | 175 (104 – 272) | 159 | 13 (12 – 13) | 105 (88 – 116) | 90 | 0.0 | 99.4 (99 - 99.7) | 98.7 |
| **KRAS** | **50955** | 1152632 (585347 – 1719916) | 8025802 (5671543 – 9971994) | 2890215 | 44 (20 – 68) | 237 (170 – 289) | 101 | 253 (218 – 287) | 251 (155 – 344) | 100 | 23 (11 – 34) | 158 (111 – 196) | 57 | 0.0 | 99.8 (99.6 - 99.9) | 28.7 |
| **BRCA2** | **91218** | 299552 (230233 – 368871) | 5052621 (3318290 – 6015311) | 3226173 | 12 (9 – 14) | 135 (86 – 170) | 115 | 41 (34 – 48) | 86 (62 – 112) | 62 | 3 (3 – 4) | 55 (36 – 66) | 35 | 0.0 | 27.8 (0.1 – 44) | 9.7 |
| **PAX7** | **120934** | 565060 (516425 – 613694) | 3748292 (2229649 – 5121684) | 2968806 | 29 (27 – 30) | 182 (83 – 253) | 142 | 62 (43 – 81) | 46 (40 – 57) | 43 | 5 (4 – 5) | 31 (18 – 42) | 25 | 0.0 | 10.3 (0 - 19.3) | 4.3 |
| **APC** | **144265** | 556426 (506785 – 606066) | 3208880 (2036481 – 4455271) | 1908177 | 23 (15 – 30) | 116 (74 – 154) | 82 | 51 (36 – 67) | 34 (28 – 44) | 23 | 4 (4 – 5) | 22 (14 – 31) | 13 | 0.0 | 3.4 (0 - 5.7) | 0 |
| **Total Target** | **564206** | 8118583 (6239144 – 9998021) | 56791795 (29425097 – 84518620) | 54704217 | 484 (354 – 613) | 2992 (1412 – 4874) | 3790 | 181 (151 – 210) | 150 (94 – 194) | 171 | 15 (12 – 19) | 106 (55 – 158) | 97 | 4.0 (2.2 – 5.7) | 37.4 (26.8 – 43) | 27.9 |
| **Total Run** | | 276025707 (175549380 – 376502034) | 2409003747 (1076062785 – 3574634209) | 1894903771 | 17281 (11798 – 22764) | 111274 (46236 – 178027) | 102727 | | | | | | | | | |

For some conditions, more than one run was performed during the testing phase. For these runs, average values are represented in the table, with ranges in parenthesis. * For one of the three MCF10A ACME runs, only 3 samples were pooled together instead of 4. All other runs had 4 samples pooled together for each run.

### Supplemental Table 3

**Supplemental Table 3. Run summaries for the targeted cancer panel genes across non-ACME and ACME single library sequencing runs**

| | | Bases | | | Reads | | | Fold enrichment | | | Coverage | | | % Bases over 60x | | |
|---|---|---|---|---|---|---|---|---|---|---|---|---|---|---|---|---|
| **Gene** | **Size** | **MCF10A No ACME** | **MCF10A ACME** | **SKBR3 ACME** | **MCF10A No ACME** | **MCF10A ACME** | **SKBR3 ACME** | **MCF10A No ACME** | **MCF10A ACME** | **SKBR3 ACME** | **MCF10A No ACME** | **MCF10A ACME** | **SKBR3 ACME** | **MCF10A No ACME** | **MCF10A ACME** | **SKBR3 ACME** |
| **MYC** | **12565** | 1125143 (761256 – 1489029) | 4036529 | 3908616 (718214 – 7099017) | 104 (73 – 134) | 367 | 361 (56 – 665) | 2039 (1279 – 2800) | 1880 | 5503 (1417 – 9590) | 90 (61 – 118) | 321 | 311 (57 – 565) | 84.9 (70.2 – 99.6) | 99.9 | 68.2 (36.3 – 100) |
| **HOXA9** | **18506** | 430837 (196385 – 665288) | 1700199 | 879713 (162145 - 1597281) | 26 (9 – 43) | 86 | 41 (4 – 77) | 439 (388 – 490) | 538 | 841 (217 – 1465) | 23 (11 – 36) | 92 | 48 (9 – 86) | 0 | 99.3 | 49.5 (0 – 98.9) |
| **FGFR4** | **19916** | 846694 (469763 – 1223625) | 2764208 | 337106 (31193 - 643018) | 58 (35 – 81) | 163 | 24 (1 – 47) | 876 (663 – 1090) | 812 | 293 (39 – 548) | 43 (24 – 61) | 139 | 17 (2 – 32) | 41.1 (0 – 82.2) | 99.8 | 0.0 |
| **STK11** | **30286** | 780738 (533973 – 1027503) | 1991085 | 314858 (85600 - 544116) | 35 (27 – 42) | 90 | 17 (4 – 30) | 590 (366 – 815) | 385 | 187 (70 – 305) | 26 (18 – 34) | 66 | 10 (3 – 18) | 0 | 94.5 | 0.0 |
| **CDKN2A** | **30774** | NA | NA | 484790 (120378 - 849201) | NA | NA | 16 (7 – 25) | NA | NA | 283 (97 – 468) | NA | NA | 16 (4 – 28) | NA | NA | 0.0 |
| **TERT** | **44787** | 475792 (409047 – 542536) | 1481668 | 486263 (115294 - 857231) | 17 (16 – 18) | 37 | 23 (7 – 38) | 276 (131 – 422) | 194 | 194 (64 – 325) | 11 (9 – 12) | 33 | 11 (3 – 19) | 0 | 0 | 0.0 |
| **KRAS** | **50955** | 731042 (560925 – 901159) | 2953982 | 634286 (238744 - 1029828) | 22 (16 – 27) | 83 | 26 (11 – 41) | 350 (191 – 509) | 339 | 230 (116 – 343) | 14 (11 – 18) | 58 | 12 (5 – 20) | 0 | 23.2 | 0.0 |
| **BRCA2** | **91218** | 326143 (321310 – 330975) | 1717813 | 282843 (201237 - 364449) | 9 (9 – 9) | 35 | 8 (7 – 9) | 101 (39 – 163) | 110 | 61 (55 – 68) | 4 (4 – 4) | 19 | 3 (2 – 4) | 0 | 0 | 0.0 |
| **PAX7** | **120934** | 402730 (357362 – 448097) | 2544000 | 532198 (191325 - 873070) | 17 (7 – 27) | 69 | 20 (9 – 31) | 88 (40 – 137) | 123 | 81 (39 – 123) | 3 (3 – 4) | 21 | 5 (2 – 7) | 0 | 0 | 0.0 |
| **APC** | **144265** | 204199 (169299 – 239099) | 942351 | 345128 (177775 - 512480) | 8 (7 – 8) | 28 | 13 (8 – 17) | 45 (13 – 77) | 38 | 45 (31 – 60) | 2 (2 – 2) | 7 | 3 (2 – 4) | 0 | 0 | 0.0 |
| **Total Target** | **533432** | 5323316 (3849120 – 6797511) | 20131835 | 8205798 (2041905 - 14369691) | 294 (202 – 386) | 958 | 547 (114 – 980) | 223 (130 – 315) | 209 | 261 (90 – 432) | 10 (7 – 13) | 38 | 15 (4 – 25) | 3.35 (1.6 – 5.1) | 16.2 | 3.2 (0.8 – 5.5) |
| **Total Run** | | 190609890 (72673305 – 308546475) | 568769190 | 165779784 (134289629 - 197269938) | 10299 (4050 – 16547) | 22818 | 9538 (7806 – 11270) | | | | | | | | | |

For some conditions, more than one run was performed during the testing phase. For these runs, average values are represented in the table, with ranges in parenthesis.

<u>Supplemental Table 4</u>

**Supplemental Table 4. Number of reads spanning X% of gene for each targeted cancer panel gene across non-ACME and ACME single library sequencing runs**

| | | MCF10A No ACME | | | | MCF10A ACME | | | | SKBR3 ACME | | | |
|---|---|---|---|---|---|---|---|---|---|---|---|---|---|
| **Gene** | **Size** | **100%** | **80%** | **60%** | **40%** | **100%** | **80%** | **60%** | **40%** | **100%** | **80%** | **60%** | **40%** |
| **MYC** | **12565** | 85 | 91 | 93 | 96 | 303 | 324 | 331 | 342 | 278 | 304 | 318 | 337 |
| **HOXA9** | **18506** | 34 | 38 | 45 | 47 | 137 | 142 | 143 | 145 | 7 | 15 | 16 | 17 |
| **FGFR4** | **19916** | 23 | 23 | 24 | 24 | 88 | 92 | 93 | 97 | 42 | 45 | 49 | 53 |
| **STK11** | **30286** | 22 | 23 | 25 | 28 | 57 | 60 | 65 | 69 | 9 | 10 | 10 | 10 |
| **CDKN2A** | **30774** | NA | NA | NA | NA | NA | NA | NA | NA | 12 | 12 | 13 | 17 |
| **TERT** | **44787** | 6 | 8 | 9 | 12 | 23 | 26 | 31 | 35 | 1 | 2 | 2 | 13 |
| **KRAS** | **50955** | 10 | 11 | 14 | 15 | 41 | 48 | 55 | 61 | 7 | 8 | 10 | 14 |
| **BRCA2** | **91218** | 1 | 2 | 3 | 4 | 7 | 9 | 12 | 20 | 1 | 2 | 2 | 4 |
| **PAX7** | **120934** | 1 | 1 | 1 | 3 | 4 | 9 | 12 | 20 | 1 | 1 | 1 | 4 |
| **APC** | **144265** | 0 | 0 | 0 | 1 | 2 | 2 | 2 | 4 | 0 | 1 | 1 | 2 |

For some conditions, more than one run was performed during the testing phase. For these runs, average values are represented in the table.
